# The mechanism of DRB7.2:DRB4 mediated sequestering of endogenous inverted-repeat dsRNA precursors in plants

**DOI:** 10.1101/2025.01.15.633165

**Authors:** Sneha Paturi, Debadutta Patra, Priti Chanda Behera, Ramdas Aute, Nilam Waghela, Priyadarshan Kinatukara, Mandar V. Deshmukh

## Abstract

Noncoding transcribable inverted repeat sequences are vital in genome stability, regulation of transposable elements, mutations, and diseases in eukaryotes. In vascular plants, dsRNA Binding Proteins (dsRBPs), DRB7.2 and DRB4, inhibit Dicer-like-protein 3 (DCL3) to stall endogenous inverted-repeat dsRNA (endo-IR dsRNA) mediated gene silencing. As dsRBPs generally assist Dicers, the inhibition of DCL3 by a dsRBP complex is quite enigmatic. Here, we explore how the DRB7.2:DRB4 complex sequesters substrate dsRNA of DCL3 using a structure-based mechanistic approach. Intriguingly, the crucial step of endo-IR dsRNA precursor sequestration is the high affinity complex formation of interacting domains of DRB7.2 (DRB7.2M) and DRB4 (DRB4D3). Next, we establish that DRB7.2 simultaneously interacts with DRB4 and endo-IR dsRNA precursors, where DRB4 contributes towards enhancement in the affinity of the complex with dsRNA, thereby impairing DCL3 mediated cleavage of endo-IR dsRNA precursors. The uniqueness of the DRB4D3 structure implies that the trans-acting (tasi)/siRNA initiation complex formed by DCL4:DRB4 in plants is diverse from its non-plant higher eukaryotes. Overall, we present considerable insights into endo-IR dsRNA precursors regulation in plants and indicate a differential evolution of RNAi initiation complexes between plants and other higher eukaryotes.

## INTRODUCTION

Organisms ranging from microsporidian parasites to humans and plants invoke elaborate gene regulation mechanisms mediated by small non-coding RNA during development as well as to combat infection and stress (González Plaza, 2020; Sunkar & Zhu, 2004). In general, Dicer depends on dsRNA Binding Proteins (dsRBPs), containing strictly two or more dsRNA binding domain (dsRBD), as the non-catalytic factors for enhancing the dsRNA cleavage activity. A few structurally and functionally well-characterized dsRBDs that exclusively interact with dsRNA and regulate the RNAi pathway include *Hs*TRBP dsRBD1 (Masliah *et al*, 2018), *Hs*TRBP dsRBD2 (Masliah *et al*, 2018), *Dm*R2D2 dsRBD1 (Yamaguchi *et al*, 2022), *Ce*RDE-4 dsRBD2 (Chiliveri & Deshmukh, 2014), *At*DRB1 dsRBD1 (Yang *et al*, 2010), and *At*DRB4 dsRBD1 (Chiliveri *et al*, 2017). On the other hand, *Hs*TRBP dsRBD3 (Wilson *et al*, 2015), *Dm*R2D2 dsRBD3 (Yamaguchi *et al*, 2022), and *Dm*LoqsPB dsRBD3 (Jouravleva *et al*, 2022) form a very stable complex with the corresponding Dicer. Nonetheless, dsRBDs that simultaneously interact with dsRNA and mediate protein-protein interaction are yet to be identified, except for R2D2 dsRBD2 (Yamaguchi *et al*, 2022) and DGCR8 dsRBDs (Partin *et al*, 2020), and hold potential for the unexplored functions of dsRBDs in biological processes.

Depending on the nature and the origin of the trigger dsRNA, plants recruit a specific subset of Dicer and corresponding dsRBPs to effect the gene regulation (Wilson & Doudna, 2013). For example, when presented with long *trans*-acting or viral dsRNA, DCL4 associates with DRB4 to produce a 21 nt siRNA duplex that further leads to AGO1/AGO7 mediated cleavage of the cognate transcript (Fukudome *et al*, 2011; Adenot *et al*, 2006). While the structure and functional mechanism of DCL4 is still unknown, DRB4 has two N-terminal dsRBDs with canonical α-β-β-β-α fold and an extended unstructured region followed by a small folded domain in the C-terminus with no known sequence or structural homology (Chiliveri *et al*, 2017). In addition, at various stages of viral infections and developmental stages in plants, DRB4 independently interacts with Hypersensitive Response to Turnip crinkle-virus (HRT), virus coat protein (vCP), DRB7.1, and DRB7.2 to perform a diverse set of functions in versatile gene regulatory mechanisms (Zhu *et al*, 2013; Tschopp *et al*, 2017; Montavon *et al*, 2017). However, additional studies are needed to decipher how DRB4 compartmentalizes the task of association with several partners.

Primary sequence searches across the plant genome indicate that DRB4 is conserved in all plants, whereas DRB7.1 and DRB7.2 are specific to vascular plants (Clavel *et al*, 2016). Furthermore, phylogenetic analysis suggests that DRB7.1 and DRB7.2 belong to a new clade of the dsRBD family (Type III) possessing the closest sequence homology with DCL4 dsRBD2 (Clavel *et al*, 2016). It was also proposed that the dsRBD of DRB7.1 and DRB7.2 may interact with both dsRNA and partner protein (Clavel *et al*, 2016). Earlier, Wuest *et al*. showed a relatively higher gene expression profile of DRB7.2 in egg cells, suggesting an essential role in the development of the female gametes (Wuest *et al*, 2010).

Amongst multiple partners, DRB4 interacts independently with DRB7.1 and DRB7.2 to sequester endogenous precursor dsRNA and prevent DCL3 from producing epigenetically activated siRNAs (easi-RNAs) (Tschopp *et al*, 2017) and endogenous inverted repeat siRNAs (endo-IR siRNAs) (Montavon *et al*, 2017), respectively. Additionally, Montavon *et. al*., showed that both single and double mutants of *drb7.2* and *drb4* accumulate a similar level of endo-IR siRNAs originating from the transcripts of locus IR71 and IR2039, signifying that the association of DRB7.2:DRB4 safeguards precursors of endo-IR siRNAs (Montavon *et al*, 2017). However, when the DRB7.2:DRB4 complex is dissociated, DCL3 generates 24 nt near-perfect self-complementarity products from these transcripts (Chan, 2008; Lu *et al*, 2006; Kasschau *et al*, 2007). Interestingly, DCL3 produced 24 nt dsRNA initiates RNA-directed DNA methylation (RdDM) that leads to *de novo* methylation across the centromeric and repetitive sequences in the genome (Erdmann & Picard, 2020). RdDM plays a crucial role in the stress response as well as development (Huettel *et al*, 2007) and is implicated in modulating various epigenetic processes, such as transgene silencing, transposon suppression, gene imprinting, etc. (Erdmann & Picard, 2020). Thus, the epigenetic modifications in the inverted repeat sequences are regulated through a delicate balance between the DRB7.2:DRB4 complex versus DCL3.

To the best of our knowledge, DRB7.2:DRB4 is a first-of-its-kind complex that inhibits Dicer from processing the trigger dsRNA instead of assisting it in the process. The antagonistic activity of DRB7.2:DRB4 on DCL3 poses a fundamental structural and mechanistic question in the RNAi pathway.

Using an integrated approach involving NMR and X-ray crystallography, here, we present the structure of the complex formed by the interaction domains of DRB7.2 and DRB4 (i.e., DRB7.2M and DRB4D3). Furthermore, we show that the complex interacts with the dsRNA using the exclusive free RNA binding site present on DRB7.2M and the two N-terminal dsRBDs of DRB4 to enhance the affinity of the DRB7.2:DRB4 for the substrate dsRNA binding. Our study provides the mechanism of the ternary complex formation and proposes a mechanism for the inhibition of DCL3 activity by DRB7.2:DRB4.

## RESULTS

### Domain architecture and interaction of DRB7.2 with DRB4

At the outset, using primary sequence analysis, we identified and defined domain boundaries in DRB7.2 and DRB4.

For DRB7.2, the program cNLS Mapper (Kosugi *et al*, 2009) predicted that residues Q20-L37 form a strong bipartite Nuclear Localization Signal (NLS) (Fig. 1A). As DRB7.2 exclusively localizes and functions in the nucleus (Montavon *et al*, 2017), having a strong N-terminal NLS affirms that DRB7.2 does not need additional co-factors for its localization. The disorder prediction analysis using PONDR for DRB7.2 yields two disordered regions (M1-S83 and Y163-V190) flanking a dsRBD domain (S84-G162) (Fig. 1A). As the N- and C-terminal disordered regions in DRB7.2 may interfere with the structural studies of the full-length protein, we additionally cloned the dsRBD region as T71-G162, marked as DRB7.2M (Fig. 1A). The N-terminal flanking residues T71-S83 provided stability to the dsRBD domain as constructs devoid of it were not stable. Additionally, we have cloned and expressed only the N-terminal region of DRB7.2 (DRB7.2N, i.e., M1-Q81) (Fig. 1A).

**Figure 1:**
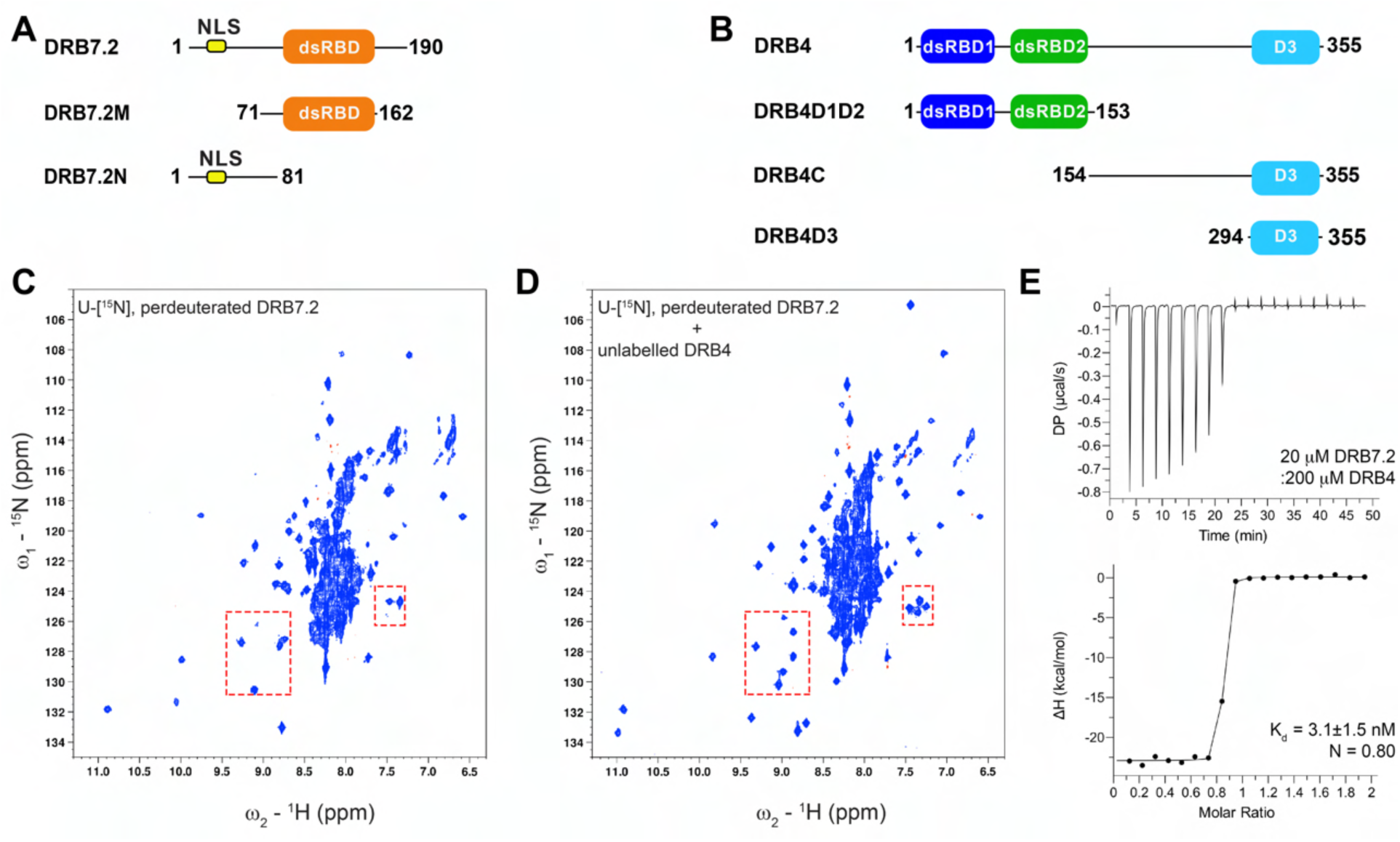
Domain architecture, characterization of DRB7.2, and its interaction with DRB4. (A) Various DRB7.2 constructs used in the study are represented as full-length, dsRBD containing region annotated as DRB7.2M (orange), and the unstructured N-terminus. The nuclear localization sequence (NLS) composed of residues Q20-L37 is marked with a yellow box. (B) Various DRB4 constructs are shown as DRB4, DRB4D1D2, DRB4C and DRB4D3. Structured domains are color coded as dsRBD1 (blue), dsRBD2 (green), and D3 (cyan). (C) ^1^H-^15^N TROSY-HSQC spectrum of perdeuterated, U-[^15^N] DRB7.2. For a ∼ 22 kDa complex, the spectral crowding between 7.5-8.5 ppm in the ^1^H dimension and overall broad resonances indicate that DRB7.2 is undergoing heterogeneous conformational exchange. (D) The ^1^H-^15^N TROSY-HSQC of DRB7.2 in the presence of equimolar DRB4. The formation of the complex shows significant improvement in the spectrum (representative red rectangles in (C and D)). (E) The isothermal calorimetric titration studies between DRB7.2 and DRB4 reveal that these two proteins form a high affinity complex. The top panel depicts the thermogram for the titration of 2 μL consecutive injections of 200 μM DRB4 into 20 μM DRB7.2. The corresponding bottom panel is the normalized binding isotherm, showing integrated changes in enthalpy (ΔH) against the molar ratio. The titration of DRB7.2 with DRB4 yielded K_d_ = 3.1±1.5 nM with N = 0.80.

Similarly, based on our previous studies (Chiliveri *et al*, 2017), we have prepared four clones of DRB4, namely, DRB4, DRB4D1D2, DRB4C, and DRB4D3 (Fig. 1B). DRB4D1D2 consists of both dsRBD domains, whereas the DRB4C is primarily disordered with a small structured domain located at the end of the C-terminal region, namely DRB4D3.

To establish the interaction between DRB7.2 and DRB4, we monitored the changes in the ^1^H– ^15^N TROSY-HSQC spectrum of DRB7.2 in the presence of the saturating concentrations of unlabeled DRB4. DRB7.2 shows many well-dispersed broad resonances, probably due to the underlying exchange processes of partially folded states or a heterogeneous ensemble of interconverting oligomeric states (Fig. 1C). Similarly, narrow dispersion in the proton dimension seen for many resonances is probably emanating from long disordered regions in DRB7.2 (Fig. 1C). Interestingly, upon addition of the molar equivalence of unlabeled DRB4 to DRB7.2, we observed significant line narrowing and the appearance of new resonances, albeit with the unchanged central region in the spectrum (Fig. 1D). Subsequent ITC studies yield an exothermic and enthalpically driven binding yielding a dissociation constant (K_d_) of 3.1 nM for the interaction between DRB7.2 and DRB4 suggesting that these proteins form a high-affinity complex by associating at an equimolar ratio as N is ∼ 0.8 (Fig. 1E and Table 1).

**Table 1:**
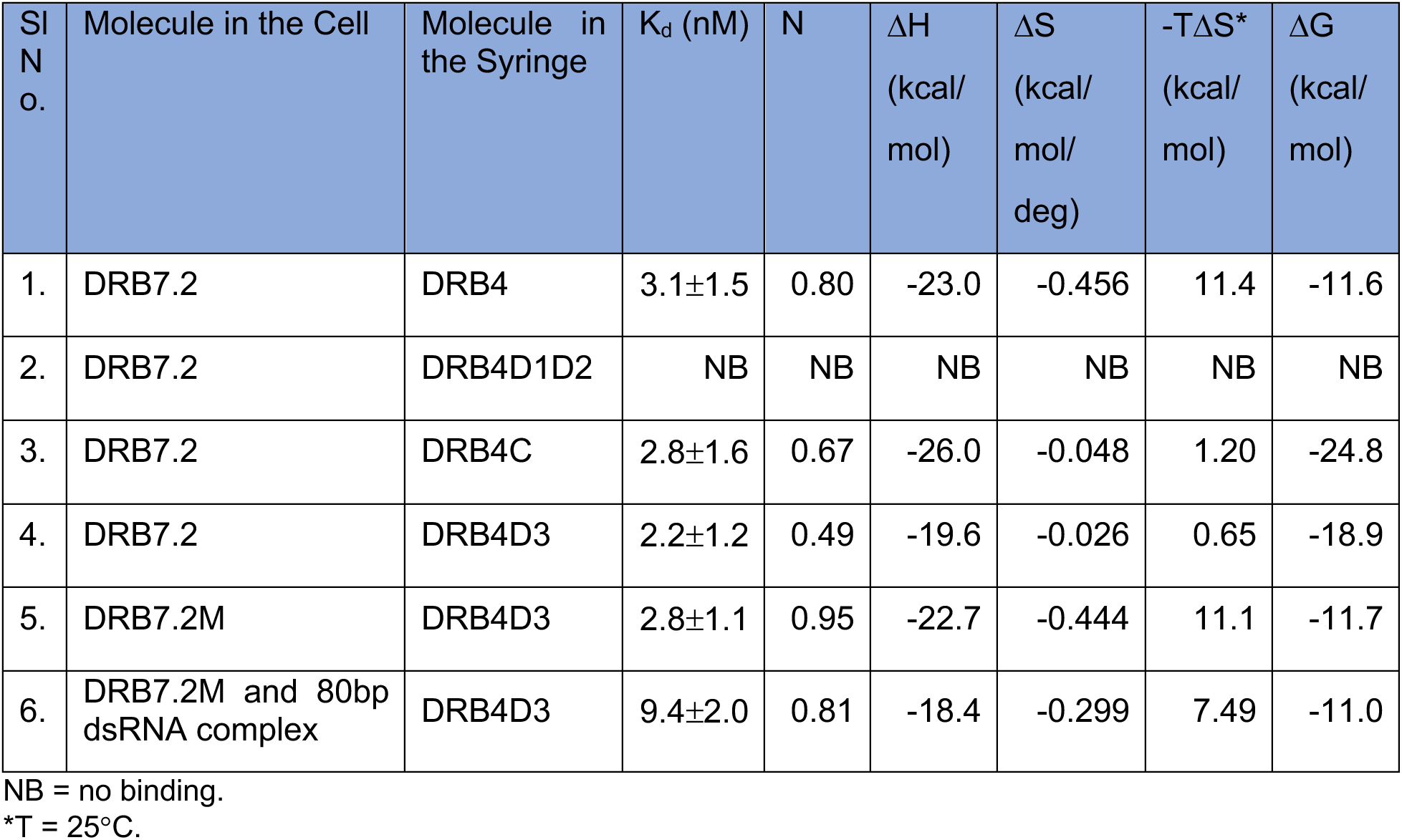
Thermodynamic parameters derived from ITC studies.

### The nature of dsRBD region of DRB7.2

To identify the interaction domains between DRB7.2 and DRB4, we performed ITC titrations of DRB7.2 with DRB4D1D2, DRB4C, and DRB4D3. While DRB4D1D2 does not bind with DRB7.2, the isotherm of DRB7.2:DRB4C yields a K_d_ ∼ 2.8 nM, and DRB7.2:DRB4D3 interactions with a K_d_ ∼ 2.2 nM (Fig. S1A-S1C and Table 1). As the thermodynamic parameters for the titrations of DRB7.2:DRB4 and DRB7.2:DRB4C are highly similar to DRB7.2:DRB4D3, we reason that DRB4D3 is solely responsible for mediating the interaction between DRB7.2 and DRB4.

Similarly, we proceeded to identify if DRB7.2 possesses a specific region that drives its interaction with DRB4. Hence, we individually titrated the unlabeled DRB4C to ^15^N-DRB7.2N and ^15^N-DRB7.2M and monitored the changes in the ^1^H-^15^N TROSY-HSQC. For DRB7.2N:DRB4C titration, we did not observe any changes in the spectral pattern of amide resonances, suggesting that the DRB4C does not bind to DRB7.2N (data not shown). At the same time, the ^1^H-^15^N TROSY-HSQC of DRB7.2M, as shown in Fig. 2A, displays hallmark features observed for DRB7.2, such as significantly broadened resonances probably arising from conformational heterogeneity or due to the presence of higher order oligomers of alone DRB7.2M. The ^15^N backbone relaxation studies on DRB7.2M also indicate a *τ*_c_ ∼ 11.89 ns based on the R_2_*R_1_ and R_2_/R_1,_ which is much higher than a theoretical *τ*_c_ ∼ 5-6 ns for a globular protein of this size (Fig S2). These data further imply that DRB7.2M is probably experiencing slow to intermediate exchange processes induced by monomer:dimer and interconverting higher-order oligomeric states in equilibrium.

**Figure 2:**
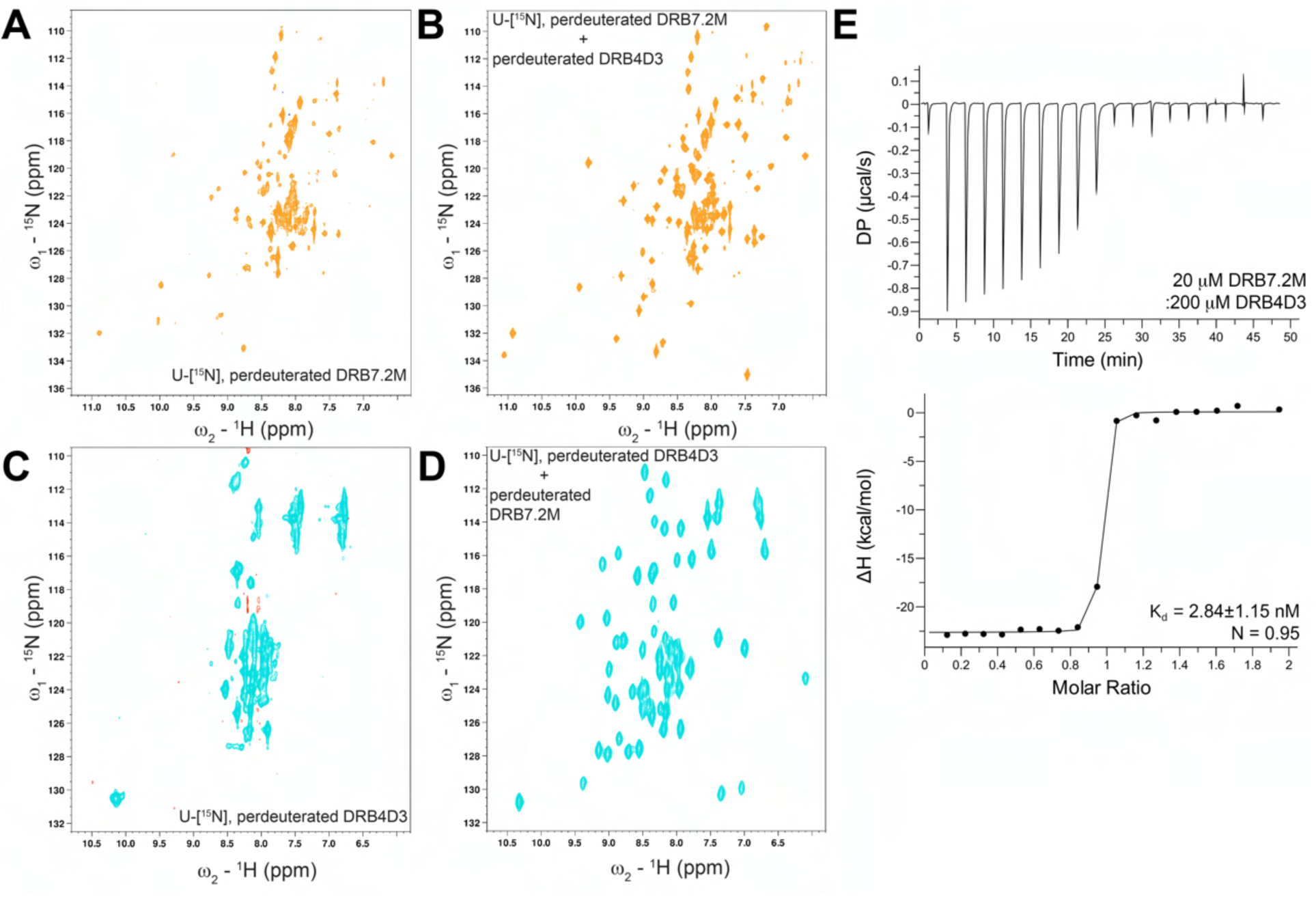
Interaction studies of DRB7.2M and DRB4D3. A ^1^H-^15^N TROSY-HSQC of (A) DRB7.2M, (B) DRB4D3, (C) equimolar complex of ^15^N DRB7.2M with unlabelled DRB4D3, and (D) equimolar complex of ^15^N DRB4D3 with unlabelled DRB7.2M. (E) The binding interaction between DRB7.2M and DRB4D3 titration is probed by ITC using 2 μL consecutive injections of 200 μM DRB4D3 into 20 μM DRB7.2M. The normalized binding isotherm fitted with K_d_ = 2.84±1.15 nM with N = 0.95 for integrated changes in enthalpy (ΔH) against the molar ratio.

Furthermore, the ^1^H-^15^N TROSY-HSQC spectra of DRB7.2M showed similar line narrowing upon titration of unlabeled DRB4D3, identical to those observed for DRB7.2, indicating that the DRB7.2M:DRB4D3 attain a well-defined complex structure (Fig. 2B). These data suggest that DRB7.2M is probably necessary and sufficient to achieve the specific interaction with DRB4.

The broad resonances in the DRB7.2M triggered us to perform size exclusion chromatography followed by MALDI-TOF measurements (Fig. S3A-S3B) to understand the nature of conformational heterogeneity in DRB7.2M. We observe that DRB7.2M (M_t_ = 10670 Da) elutes at an MW 10440 Da (SEC) and gives a major MALDI peak corresponding to the monomeric mass at 12009 Da (∼ 85%) and a small dimeric population at 24184 Da (∼ 15%).

Previously, using phylogenetic analysis, Clavel *et al*., showed that DRB7.2 dsRBD belongs to a type III class of the dsRBD family and harbors a close relationship with *At*DCL4 dsRBD2 (Clavel *et al*, 2016). Our primary sequence analysis of DRB7.2M with dsRBDs of *Hs*TRBP, *Dm*R2D2, and *Ce*RDE-4 suggests that the dsRBD fold is well conserved in DRB7.2M (Fig. S3C).

### DRB7.2M and DRB4D3 constitute interaction regions in DRB7.2:DRB4 complex

Our earlier studies showed that DRB4C is comprised of two dominant regions consisting of a highly flexible segment composed of N154-S293 and a domain encompassing E294-P355 (DRB4D3) (Chiliveri *et al*, 2017). As seen in Fig. 2C, ^1^H-^15^N TROSY-HSQC of DRB4D3 shows fewer than expected peaks that have a significantly narrow dispersion and larger line widths. Moreover, DRB4D3 (M_t_ = 7810 Da) elutes at a higher MW of 13319 Da on SEC (Fig. S3D) but shows a monomeric population at 8128 Da (∼ 85%), a dimer at 16400 Da (∼ 11%), a trimer at 24686 Da (∼ 3%), and a tetramer at 32955 Da (∼ 1%) on MALDI (Fig. S3E). These observations imply that DRB4D3 experiences a high hydrodynamic radius arising from the formation of higher-order oligomers coupled with various conformational exchange processes.

Despite extensive bioinformatics searches, DRB4D3 did not show sequence homology with any protein or domain in the UniProt database. As the homolog of DRB4 in *H. sapiens* (TRBP) and *D. melanogaster* (R2D2, Loqs-PB) also contain a C-terminal third domain tailored for an association with Dicer, we carried out the alignment of DRB4D3 with the TRBPD3, R2D2D3, and LoqsD3 (Fig. S3F). However, DRB4D3 bears only 6.4% identity and 11.9% similarity with *Hs*TRBPD3, indicating that DRB4D3 does not have any evolutionary relationship with domains that are known to interact with respective Dicers.

To know if DRB7.2M and DRB4D3 drive the specific interaction necessary to form the DRB7.2:DRB4 complex, we titrated DRB7.2M with DRB4D3 and *vice versa* (Fig. 2B and 2D). The ^1^H-^15^N TROSY-HSQC spectrum of both domains in complex form yields a highly dispersed spectrum as well as a narrow line width for most resonances. The significant changes in spectral pattern in Fig. 2B and 2D imply that DRB7.2M and DRB4D3 attain a stable conformation as a complex. The changes seen in these spectra also corroborate with the spectrum of the DRB7.2:DRB4 complex (*vide supra*). Furthermore, the ITC studies on DRB7.2M:DRB4D3 follow an identical binding pattern as for DRB7.2:DRB4 titration, which most likely arises from the folding into a stable monomeric conformation of DRB7.2M and DRB4D3 (Fig. 2E). The single-site binding isotherm results in a K_d_ ∼ 2.84 nM with approximately 1:1 stoichiometry. We also found that the DRB7.2M:DRB4D3 complex (M_t_ = 18480 Da) shows MW of 27658 Da on SEC and a corresponding equimolar complex peak at 20290 Da on MALDI (Fig. S3G-S3H). The additional peaks seen in this MALDI correspond to the fragments seen in the individual MALDI profiles of DRB7.2M and DRB4D3.

Collectively, DRB7.2M and DRB4D3 are necessary and sufficient to facilitate the binding between DRB7.2:DRB4 as they are the interaction domains. Previously, it was observed that the dsRBPs in humans and flies (e.g., TRBP, R2D2, Loqs-PB) interact with corresponding Dicers through a C-terminal domain, termed as D3, attain a dsRBD fold. If DRB4D3 possesses a putative role in DCL4 recognition, similar to TRBPD3, R2D2D3, and Loqs-PBD3, it remains enigmatic at this stage. Therefore, the evolution in plants to select DRB4D3 for an association with DRB7.2 poses an interesting conundrum to the hypothesis of DCL4:DRB4 association in the tasi/siRNA pathway.

### Solution structure of DRB7.2M in complex with unlabeled DRB4D3

To understand the molecular basis of the DRB7.2:DRB4 complex formation, we have performed structural and biochemical characterization on the DRB7.2M and DRB4D3.

As the standalone DRB7.2M and DRB4D3 are not amenable for structural studies due to significant spectral broadening (Fig. 2A and 2C), we have isotopically enriched one component of the complex while keeping the other perdeuterated to obtain better signal-to-noise ratio (Gardner & Kay, 1998). For the solution structure determination of DRB7.2M, we have used a complex formed by ^15^N, ^13^C, perdeuterated DRB7.2M with perdeuterated DRB4D3, whereas for the structure determination of DRB4D3, a complex formed by ^15^N, ^13^C, perdeuterated DRB4D3 with perdeuterated DRB7.2M is used.

The solution structure of DRB7.2M suggests that the dsRBD region (A85-V161) adopts a canonical α1-β1-β2-β3-α2 fold, in which two α helices are packed against a three-stranded anti-parallel β sheet (Fig. 3A). Details of the NMR restraints and statistics are summarized in Table 2. The ensemble of the ten lowest energy structures converged with a backbone RMSD of 0.82 Å for the dsRBD region (Fig. 3A).

**Figure 3:**
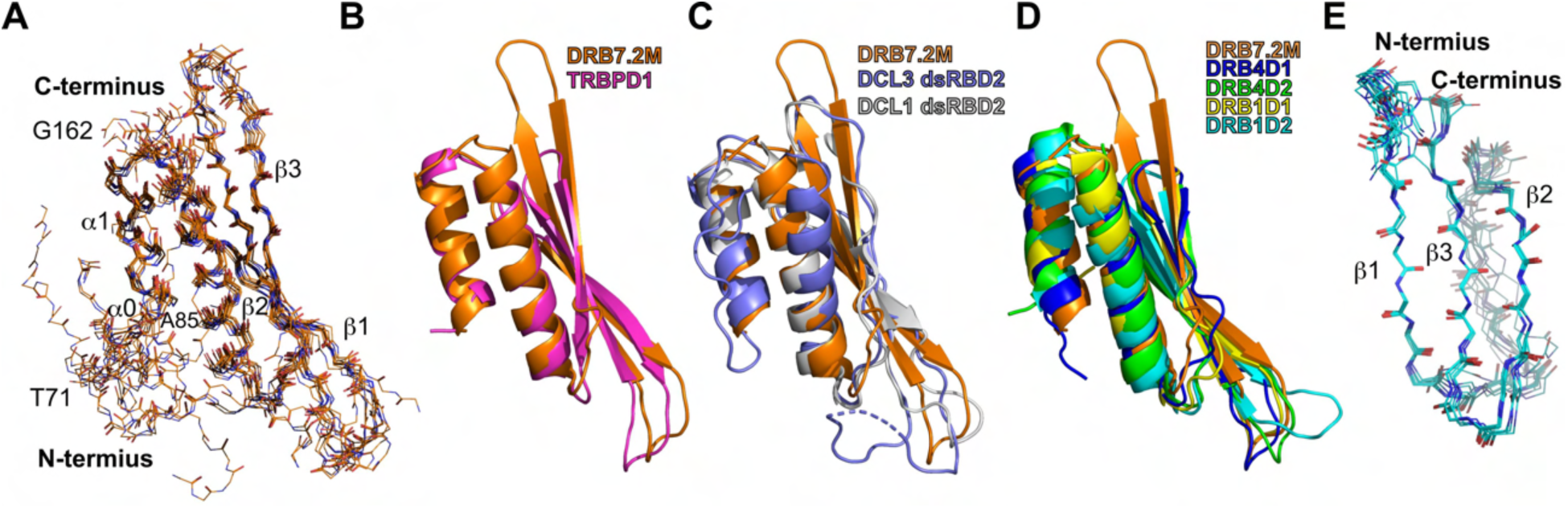
Solution structure of DRB7.2M and DRB4D3. (A) A superposition of the 10 lowest energy ensemble structures of DRB7.2M (average backbone RMSD = 0.82 Å) shows a highly conserved secondary structure in the dsRBD region. The N-terminal region (T71-S83) adopts random orientation to the DRB7.2 dsRBD. Color representation: carbon: orange; oxygen: red; nitrogen: blue. Residues A85, L89, V120, V122, L133, A150, A151, A154, and L155 form the core of DRB7.2 dsRBD. (B) The structural overlay of DRB7.2M with *Hs*TRBPD1 (PDB ID: 5N8M) yields a backbone RMSD of 1.16 Å. While most of the dsRBD region of DRB7.2 aligns well with the TRBP dsRBD1, the orientation of the β1-β2 loop is different, and DRB7.2M has a longer β2-β3 loop. (C) Overlay of DRB7.2M with DCL3 dsRBD (1.69 Å, PDB ID: 7VG2) and DCL1 dsRBD2 (1.17Å, PDB ID: 2LRS). (D) Overlay of DRB7.2M with other structured dsRBDs involved in the plant RNAi pathway, such as DRB4D1 (0.9 Å, PDB ID: 2N3G), DRB4D2 (1.9 Å, PDB ID: 2N3H), DRB1D1 (1.0 Å, PDB ID: 2L2N) and DRB1D2 (1.7 Å, PDB ID: 2L2M). The alignment shows the overall conservation of the dsRBD fold in DRB7.2 except for the extended DRB7.2 specific β2-β3 loop. (E) Ensemble of 10 lowest energy structures of DRB4D3 calculated using Rosetta with backbone RMSD 0.25 Å. β1: V323-P327, β2: A338-R343, and β3: F347-L352, whereas, V324, Y350, I348, R326 A349, F342, E339, R351, A338, P335 form hydrogen bonds, and F347, P327, W328, P330, V324, and L352 make hydrophobic contacts. Although the solution structures of DRB7.2M and DRB4D3 represented above appear as solo domains, the data was collected on the complex with an unlabeled corresponding partner, which is invisible.

**Table 2:**
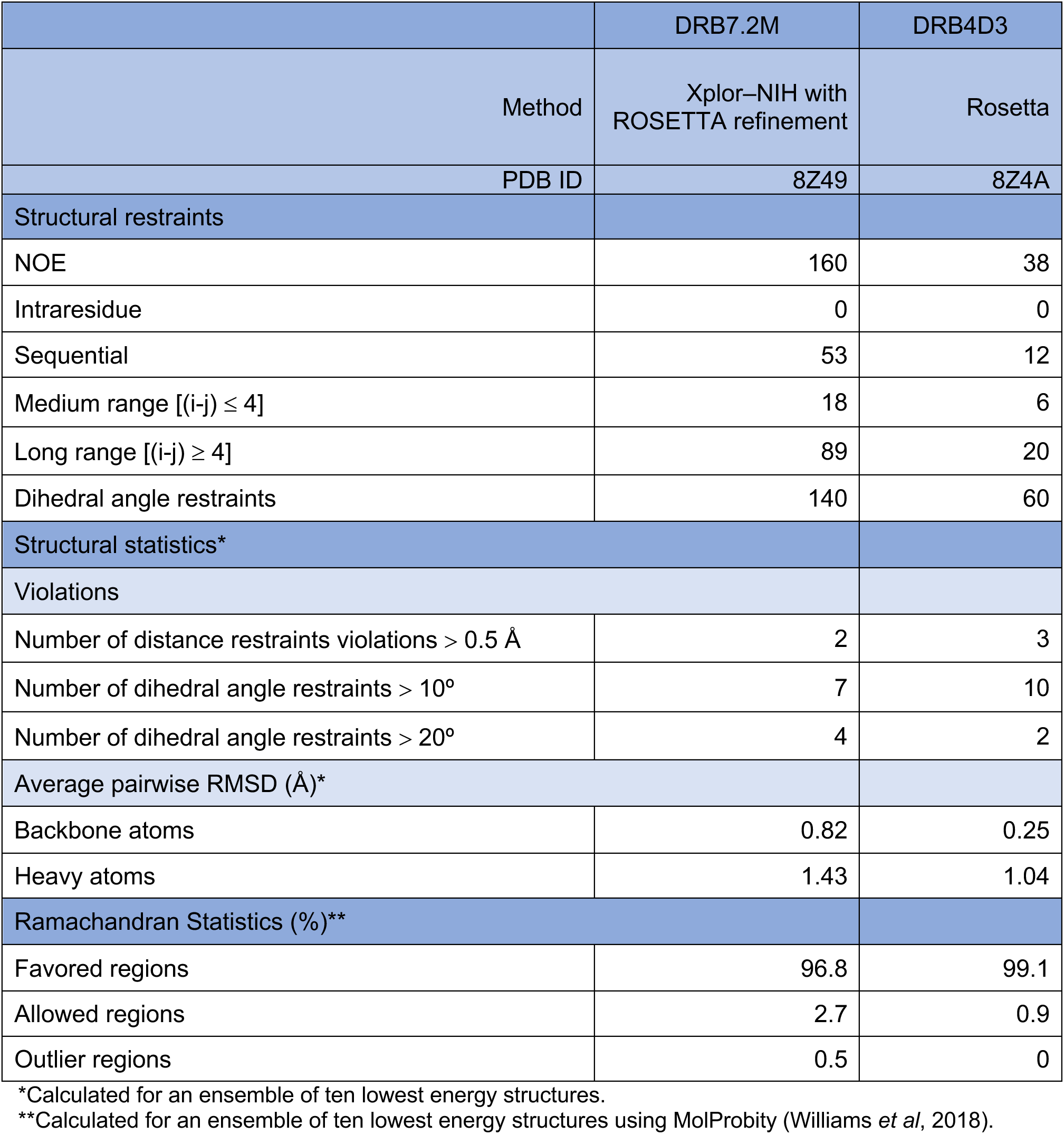
Structural statistics for the solution structure of DRB7.2M and DRB4D3.

|  | DRB7.2M | DRB4D3 |
| --- | --- | --- |
| Method | Xplor–NIH with<br>ROSETTA refinement | Rosetta |
| PDB ID | 8Z49 | 8Z4A |
| Structural restraints |  |  |
| NOE | 160 | 38 |
| Intraresidue | 0 | 0 |
| Sequential | 53 | 12 |
| Medium range [(i-j) ≤ 4] | 18 | 6 |
| Long range [(i-j) ≥ 4] | 89 | 20 |
| Dihedral angle restraints | 140 | 60 |
| Structural statistics* |  |  |
| Violations |  |  |
| Number of distance restraints violations > 0.5 Å | 2 | 3 |
| Number of dihedral angle restraints > 10° | 7 | 10 |
| Number of dihedral angle restraints > 20° | 4 | 2 |
| Average pairwise RMSD (Å)* |  |  |
| Backbone atoms | 0.82 | 0.25 |
| Heavy atoms | 1.43 | 1.04 |
| Ramachandran Statistics (%)** |  |  |
| Favored regions | 96.8 | 99.1 |
| Allowed regions | 2.7 | 0.9 |
| Outlier regions | 0.5 | 0 |

\*Calculated for an ensemble of ten lowest energy structures.
\*\*Calculated for an ensemble of ten lowest energy structures using MolProbity (Williams *et al*, 2018).

In the majority of dsRBDs, the β2-β3 turn is composed of ∼ 5 amino acids; however, in DRB7.2 dsRBD, it appears as a long loop involving nine amino acids. Moreover, the stretch formed by T71-S84 harbors a short α−helix, α0, encompassing E78-S83. We could not identify any specific contacts between α0 and the rest of the dsRBD fold, probably due to the lack of long-range NOEs attributed to the flexible nature of the stretch. At the same time, transient interactions between the stretch and the dsRBD cannot be ruled out as the stretch is essential in stabilizing the dsRBD. Previously, extensions that impart stability to dsRBD have been identified in other dsRBPs, such as RDE-4 and TRBP (Chiliveri & Deshmukh, 2014; Masliah *et al*, 2018). The presence of a longer β2-β3 sheet and the loop connecting both strands appears to be specific to DRB7.2 dsRBD, and such a modification has not yet been identified in any other dsRBD.

The superposition of the DRB7.2 dsRBD with its human homolog TRBP dsRBD1 (TRBPD1) shows a highly similar arrangement of the secondary structural elements with a backbone RMSD of 1.16 Å (Fig. 3B). Furthermore, the comparison of DRB7.2M with dsRBD2 of *A. thaliana* DCL1 (PDB ID: 2LRS) and *A. thaliana* DCL3 (PDB ID: 7VG2), which possess a relatively longer β2-β3 loop compared to the canonical dsRBDs but still shorter than DRB7.2M, showed near identical β2-β3 loop orientation and a backbone RMSD 1.17 Å and 1.69 Å, respectively (Fig. 3C). Similarly, the overlay of DRB7.2 dsRBD with DRB1 and DRB4 dsRBDs, which are structurally well-studied dsRBPs in *A. thaliana*, reveals that DRB7.2 dsRBD is well conserved. For these, the backbone of DRB7.2 dsRBD aligns with DRB1 dsRBD1 (DRB1D1) and DRB1 dsRBD2 (DRB1D2) with an RMSD of 1.0 Å and 1.7 Å, respectively; and DRB4 dsRBD1 (DRB4D1) and DRB4 dsRBD2 (DRB4D2) with an RMSD of 0.9 Å and 1.9 Å, respectively (Fig. 3D).

Overall, the DRB7.2 dsRBD appears to be highly conserved in features that contribute to the dsRNA recognition, except for the DRB7.2M specific β2-β3 region, which assumes an extended and stable structure.

### Solution structure of DRB4D3 in complex with unlabeled DRB7.2M

Using a similar strategy of isotope labeling and NMR methods, we have determined the solution structure of DRB4D3 with chemical shift information and sparse NOE restraints using Rosetta (Raman *et al*, 2010) (Table 2 and Fig. S4A). The chemical shift deviation from the random coil value for the backbone residues predicts that the region comprising E294-S316 is flexible, hence, it was eliminated from the structure calculation. The ten lowest energy ensemble structures of DRB4D3 show a backbone RMSD of 0.25 Å (Fig. 3E). In addition, we observe three cross peaks at lower ^15^N frequencies corresponding to 84 ppm, 84.4 ppm and 85 ppm in the ^1^H-^15^N HSQC spectrum of DRB4D3 suggesting the formation of the salt-bridge within DRB4D3 and between DRB4D3 and DRB7.2M (Fig. S5).

The structure reveals a novel arrangement of three anti-parallel β-strands as β1-β3-β2 in the three-dimensional space (Fig. S4B). Besides, the DRB4D3 assumes a stable conformational state only upon binding with the partner; therefore, the absence of such a fold in the structural database is not surprising. In the RNAi pathway, generally, the C-terminal region of most dsRBPs often has a third dsRBD, which is essential to make contact with the respective Dicer (Jouravleva *et al* 2022; Liu *et al* 2018; Wang *et al* 2021; Wilson *et al* 2015; Yamaguchi *et al* 2022). However, the third domain in DRB4 appears to have evolved into a completely novel fold, implying that DRB4 may involve a different strategy to recognize DCL4 in the tasi/siRNA pathway.

### The structure of the DRB7.2M:DRB4D3 complex

To determine the interaction interface in the complex, we crystallized DRB7.2M:DRB4D3. Well-diffracting crystals for the complex were readily obtained, but neither DRB7.2M nor DRB4D3 alone yielded crystals in crystallization screens. Since the solution structures of DRB7.2M and DRB4D3 were derived using sparse NMR restraints, structural models predicted by AlphaFold (Jumper et al, 2021) served as the template to solve the crystal structure of the complex using molecular replacement (Liebschner et al, 2019). The data collection and refinement statistics of the complex at 2.9 Å resolution are given in Table 3.

**Table 3:**
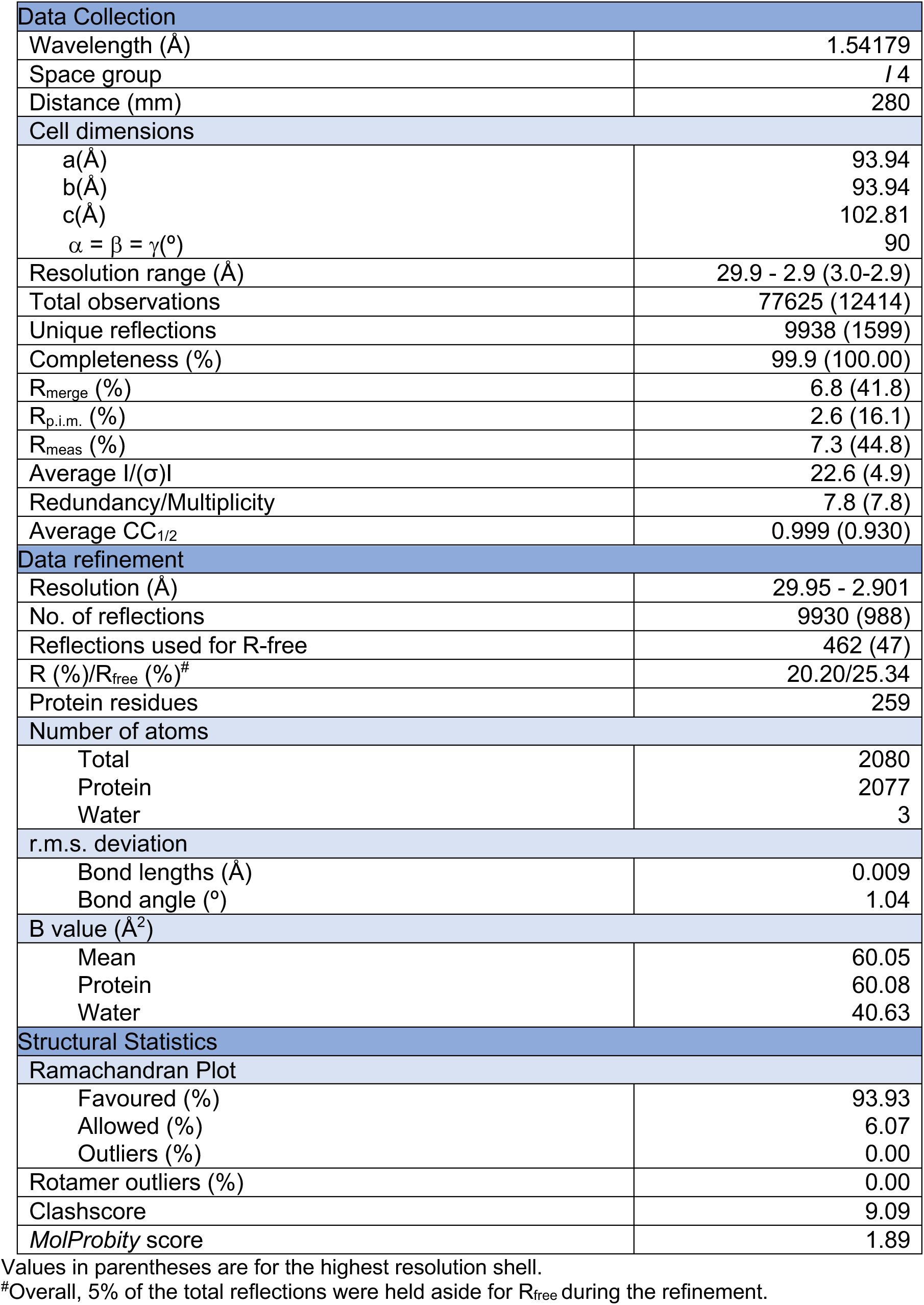
Crystallographic data collection and refinement statistics (8IGD)

The cell content analysis showed the presence of two protomers, a heterodimer of the DRB7.2M:DRB4D3, in the asymmetric unit. The complex is primarily stabilized by the β3 of DRB7.2M and β1 of DRB4D3 by burying ∼ 20% (1750 Å^2^) of the total accessible surface area (8500 Å^2^) of DRB4D3, signifying strong interaction between DRB7.2M and DRB4D3 (Fig. 4A). Besides, we find an additional interaction between the β1 strand of the two molecules of DRB7.2M in the asymmetric unit formed by burying ∼ 10% of the total surface area. However, the formation of a dimer of the heterodimeric complex appears to have been formed due to crystal packing effects. Moreover, NMR-derived ^15^N linewidths indicate that an *in vitro* reconstituted complex contains a 1:1 stoichiometry of DRB7.2M and DRB4D3.

**Figure 4:**
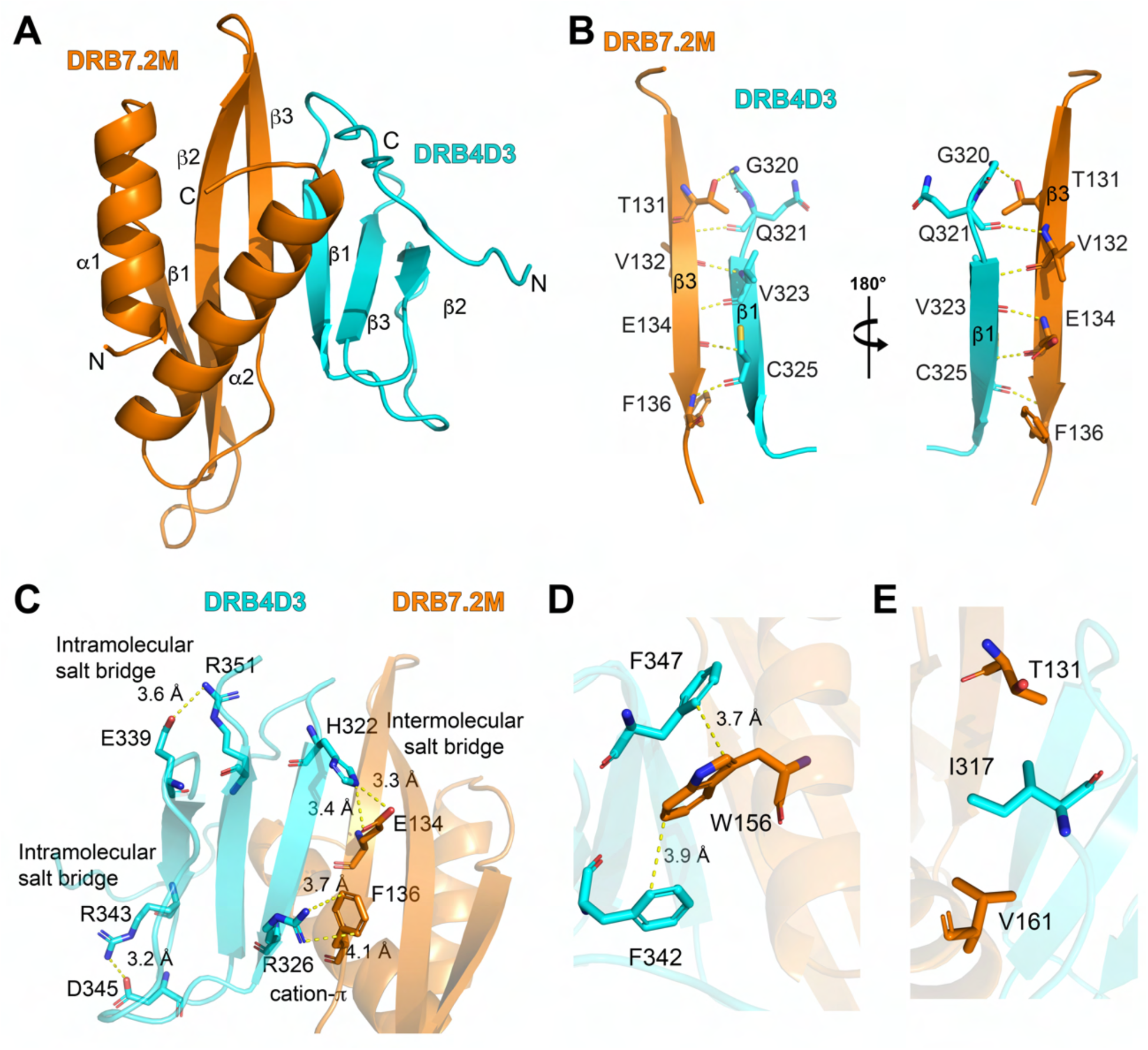
Crystal structure of DRB7.2M:DRB4D3 complex at 2.9 Å resolution. (A) The cartoon representation of the heterodimeric complex shows the helical face of DRB7.2M. In the complex, DRB7.2M (orange) adopts a canonical α1-β1-β2-β3-α2 fold, and DRB4D3 (cyan) assumes a β1-β3-β2 topology. The interaction is primarily mediated by the β3 (DRB7.2M) and β1 (DRB4D3) parallel strands. The crystal structure is solved for the regions S83-G162 of DRB7.2M and K306-P355 of DRB4D3, which showed unambiguous electron density, whereas, residues T71-R82 of DRB7.2M and E294-K305 of DRB4D3 were not visible. (B) Residues, represented as sticks and annotated, are involved in the hydrogen bond network formation between β3 (DRB7.2M) and β1 (DRB4D3). The β-sheet formation across two domains stabilizes the complex. (C) The representation of additional stabilization interactions at the binding interface between DRB7.2M and DRB4D3, such as intermolecular/intramolecular salt bridges and cation-*π* interactions, (D) Weak *π*-*π* interaction, and (E) Hydrophobic interaction. Sidechains (as a stick model) of the residues involved in the specific interactions are annotated along with the corresponding bond distances (yellow dashed lines). Atoms are represented using the color coding as oxygen (red); nitrogen (blue); and sulphur (yellow).

To further validate 1:1 stoichiometry of DRB7.2M and DRB4D3, we explored the backbone dynamics of the complex at the ps-ns timescale using R_1_, R_2_, and ^15^N {^1^H} heteronuclear NOE experiments at a 600 MHz ^1^H frequency (Fig. S6-S7). The R_1_, R_2_, and hetNOE values of DRB7.2M and DRB4D3 suggest the structured region of the complex acts as a rigid entity with no significant ps-ns time scale motions. Furthermore, the extended model-free analysis (Mandel *et al*, 1995) converged to an axially symmetric diffusion tensor with *τ*_c_ equivalent to 15.8 ns, which corresponds to the molecular weight of ∼ 24 kDa for the DRB7.2M:DRB4D3 complex (Table S1) corroborating to 1:1 stoichiometry of the complex. Besides, the hydrodynamic calculations for the dimer of the heterodimeric complex yield a theoretical *τ*_c_∼ 27 ns, which is significantly larger than the experimentally derived *τ*_c_ of the complex in the solution.

These observations conclude that the biologically functional unit is a heterodimeric complex of DRB7.2M:DRB4D3, while the interactions between the heterodimers are a product of crystal packing.

### The interaction interface between DRB7.2M:DRB4D3 complex

The structure of the complex indicates that the DRB7.2 dsRBD primarily utilizes the β-sheet region to form an interaction interface with DRB4D3 in which the y-shaped α-helical face is completely excluded. Besides, the putative dsRNA recognition residues forming the tripartite contact, such as K86, L89, and N91 in α1 helix, the tandem K143, K144 at the N-terminal α2 helix, and the invariant H112 in the β1-β2 loop appear on the surface. Moreover, the longer β3 strand in DRB7.2 dsRBD observed in the solution and crystal structure is involved in stabilizing the interaction interface of the complex (Fig. 4A). The crystal structure of DRB7.2M:DRB4D3 reveals that V132, E134, and, F136 of the β3 strands of DRB7.2M, and V323 and C325 of the β1 strand of DRB4D3 stabilize the dimerization interface through hydrogen bonding as well as polar contact formed by T131 (DRB7.2M) with G320 (DRB4D3) (Fig. 4B). Moreover, the crystal and solution structures of individual DRB7.2M and DRB4D3 are in good agreement as we observe the backbone RMSD 0.87 Å and 0.59 Å, respectively (Fig. S8A-S8B).

Further analysis reveals that the complex is stabilized by an intermolecular salt bridge formed between H322 (DRB4D3) and E134 (DRB7.2M) and a cation-*π* interaction by R326 (DRB4D3) and F136 (DRB7.2M) (Fig. 4C). Additionally, two intramolecular salt-bridges, R343:D345 and E339:R351 stabilize DRB4D3 fold in the complex (Fig. 4C). The association is also established by the C-terminus of the α2-helix of DRB7.2M with DRB4D3 through a weak *π* - *π* interactions formed by W156 of DRB7.2 with F342 and F347 of DRB4D3 (Fig. 4D) and hydrophobic interaction between V161 (DRB7.2M) and I317 (DRB4D3) (Fig. 4E). The presence of all possible non-covalent interactions mediating the association of DRB7.2M:DRB4D3 rationalizes their engagement with each other with a nano-molar affinity.

### dsRNA binding studies of the DRB7.2M:DRB4D3 complex

As the dsRNA binding surface of DRB7.2M dsRBD is available for interaction with the substrate dsRNA in the DRB7.2:DRB4D3 complex, we next asked if the DRB7.2M dsRBD forms the canonical tripartite contact with the substrate dsRNA.

In our experience, the NMR study of longer dsRNA (such as 20 bp, 40 bp or longer) with dsRBD results in the resonance broadening resulting from multiple simultaneous events, i.e., binding of multiple dsRBDs (about two to four) with a single dsRNA molecule, the intermediate scale binding affinity (such as μM affinity), the sliding of dsRBD (diffusion) over the length of dsRNA. Therefore, we titrated the DRB7.2M:DRB4D3 complex with the increasing concentration of 13 bp dsRNA, which is necessary and sufficient to interact with just one dsRBD at a time and does not lead to any deleterious effect on the ^1^H-^15^N HSQC (Fig. 5A, 5B, and S9A). Subsequently, we mapped the changes induced by 13 bp dsRNA in the resonance of ^1^H-^15^N HSQC of DRB7.2M in the complex and the normalized chemical shift perturbations were plotted as the function of residue number (Fig. 5C). The plot indicated that the residues S84, A85, K86, Q88 from α1, M113, K114, I115, F116 from β1-β2 loop, and Y141, K142, K144, A146, and E148 belonging to α2 show the major changes in the chemical environment (Fig. 5C). The NMR interaction study presented here also suggest that the integrity of DRB7.2M:DRB4D3 complex is not affected by the addition of dsRNA to the complex.

**Figure 5:**
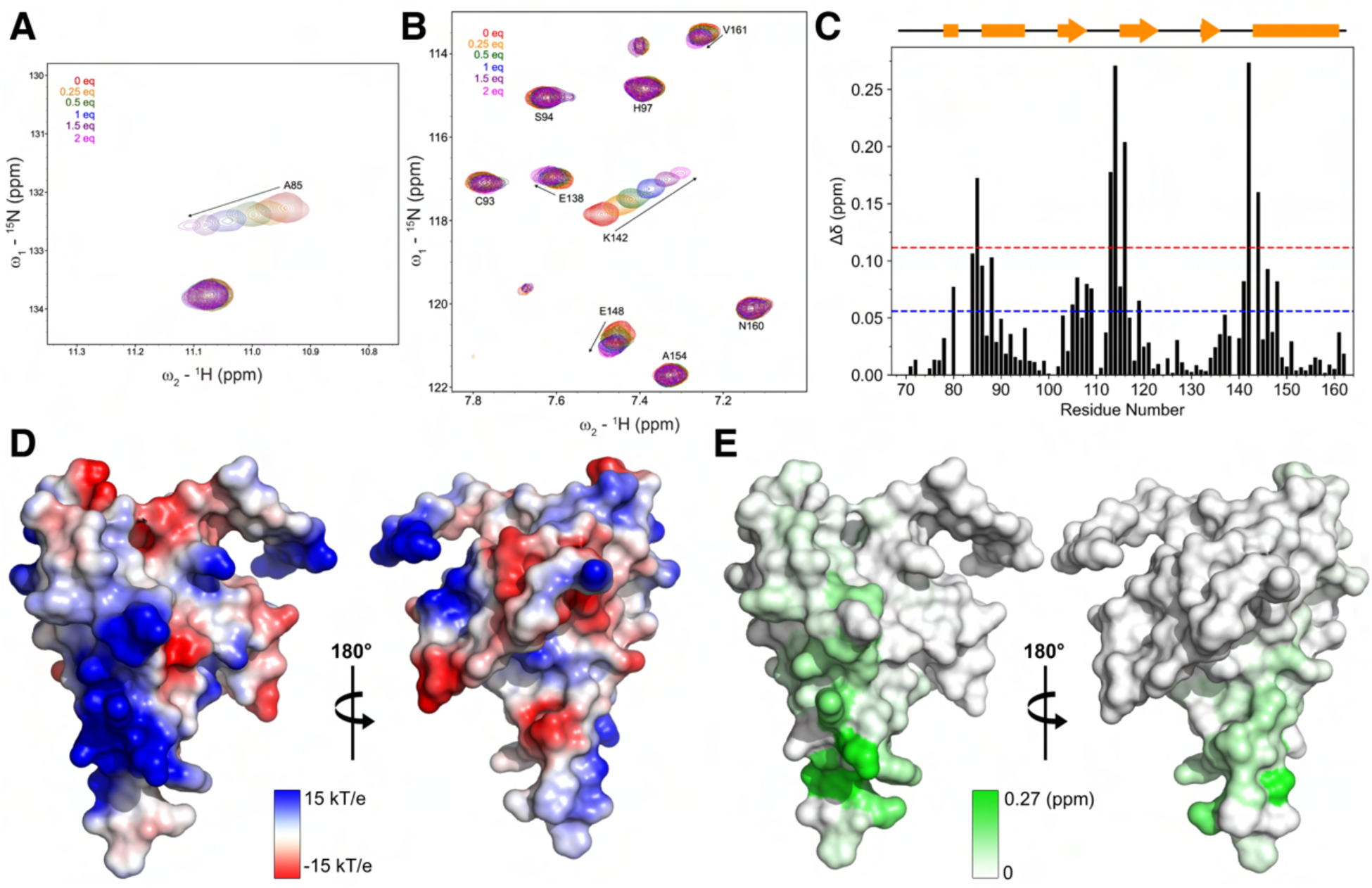
NMR-based binding studies of DRB7.2M:DRB4D3 complex with 13 bp dsRNA. (A-B) Excerpts of the chemical shift perturbation plots for the 150 μM DRB7.2M:DRB4D3 complex upon titration with various molar equivalence concentrations of 13 bp dsRNA are represented. Arrows indicate the direction of changes in the annotated residues. (C) The normalized chemical shift perturbation plot as a function of residue number indicates changes in the chemical shifts at 1:2 molar equivalence of DRB7.2M:DRB4D3 (150 μM) and 13 bp dsRNA (300 μM). The secondary structure is marked with arrows for β-sheet and rectangle for α-helical regions. The dotted lines represent statistical significance at 1α and 2α. (D) Electrostatic potential map of the surface-filled complex structure of DRB7.2M:DRB4D3 shows a contiguous positively charged patch on DRB7.2M on the α-helical face, whereas DRB4D3 shows random charge distribution. The surface charge map at −15 kT/e (red) to +15 kT/e (blue) was calculated using the Adaptive Poisson-Boltzmann Solver (APBS) Tool 2.1 plugin (Jurrus *et al*, 2018) in Pymol. (E) The surface plot of DRB7.2M:DRB4D3 complex annotated with the normalized chemical shift perturbation derived in panel C indicates that the 13 bp dsRNA induces changes in the chemical shift changes in the canonical residues arising from α1 helix, β1-β2 loop, and N-terminus of α2 helix.

As expected, the titration of DRB7.2M:DRB4D3 complex with longer dsRNA, such as 20 bp and 40 bp, led to significant resonance broadening, most likely arising due to the aforementioned reasons (Fig. S9B and S9C).

The surface electrostatic potential of the DRB7.2M:DRB4D3 shows the presence of a contiguous positive electrostatic potential on the DRB7.2M surface, whereas DRB4D3 primarily shows random charge distribution (Fig. 5D). Moreover, the positively charged surface-exposed residues in DRB7.2M are seen participating in the 13 bp dsRNA binding using the tripartite contact formed by the α1, β1-β2 loop and the N-terminus of α2 helix (Fig. 5E). The location of the dsRNA binding site lies on the opposite side of the DRB7.2M:DRB4D3 interface thus giving accessibility of dsRNA to DRB7.2M independent of DRB7.2M’s interaction with DRB4D3.

Collectively, DRB7.2M appears to engage two distinct substrates, DRB4D3 and dsRNA, with two exclusive surfaces.

### DCL3 substrate recognition by DRB7.2, DRB4, and DRB7.2:DRB4 complex

Recent cryo-EM structure of DCL3:pre-siRNA complex in an active dicing-competent state reveals that a 40 bp RNA with 5’P guide and 1 nt 3’ overhang containing complementary strand (pre-siRNA or 40 bp RNA) are recognized by PAZ:platform and connector region (Wang *et al*, 2021). Therefore, to know whether DRB7.2:DRB4 binds to the DCL3 substrate, we characterized interactions of DRB7.2, DRB4, and DRB7.2:DRB4 complex with the pre-siRNA using gel mobility shift assay.

We found that the construct bearing DRB7.2 dsRBD region binds to the 40 bp RNA (i.e., DRB7.2 and DRB7.2M), confirming that the dsRBD region in DRB7.2 is the sole dsRNA recognizer. As expected, DRB7.2N and DRB4D3 did not show any binding to the 40 bp RNA (Fig. 6A). We also found that DRB4D3 is devoid of dsRNA binding activity even at higher concentrations irrespective of the length of the dsRNA (Fig. S10A-B), which supports the earlier observation of its lack of structural features and unavailability of necessary electrostatic surface needed for dsRNA binding.

**Figure 6:**
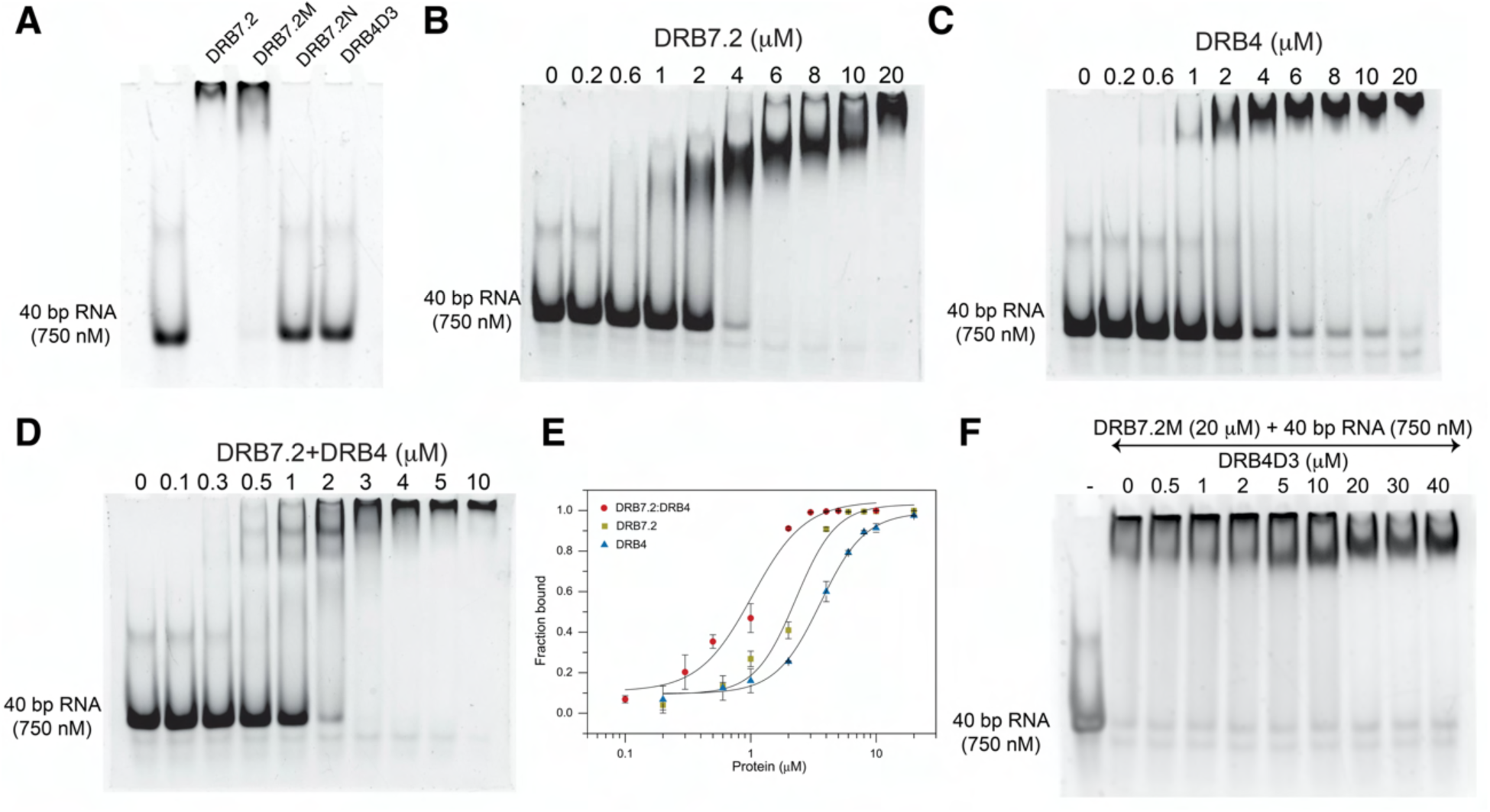
Interaction studies of DRB7.2:DRB4 complex with DCL3 dsRNA substrate. (A) Gel shift assay of 20 μM DRB7.2, DRB7.2M, DRB7.2N, and DRB4D3 with 750 nM 40 bp RNA indicating that the DRB7.2M is the sole dsRNA recognizer in DRB7.2. (B-D) Gel shift assay of 750 nM 40 bp RNA with increasing concentration of DRB7.2, DRB4, and DRB7.2:DRB4 complex. (E) Non-linear curve fit analysis to estimate the apparent K_d_ between 40 bp RNA with DRB7.2, DRB4, and their complex. DRB7.2:DRB4 shows improved affinity to the 40 bp DCL3 dsRNA substrate against the affinities exhibited by individual DRB7.2 and DRB4. (F) Gel shift assay of a preformed complex of DRB7.2M:40 bp RNA with increasing concentrations of DRB4D3 implying that the DRB7.2M:dsRNA complex is unperturbed by the DRB7.2M:DRB4D3 binding. The gels in (A-D and F) were visualized using SYBR-Gold nucleic acid gel stain.

The concentration dependent gel mobility shift assay of DRB7.2 with 40 bp RNA shows a complete saturation at 20 μM protein concentration, where distinct mobility shift intermediates are seen at protein concentrations ranging from 1 μM to 10 μM (Fig. 6B). Similarly, the concentration dependent gel mobility shift assay of DRB4 with 40 bp RNA show that DRB4 saturates around 20 μM protein concentration (Fig. 6C). In this case, we do not observe distinct mobility shift intermediates most likely due to dsRNA interaction driven by “all or none” binding possibly occurring due to the presence of two dsRBDs in DRB4. The saturation of DRB7.2:DRB4 complex with 40 bp RNA occurs by 3-4 μM protein concentration, indicating that the complex may have a relatively higher affinity of the dsRNA (Fig. 6D). Additional gel mobility shift assay studies with DRB7.2, DRB7.2M and DRB7.2M:DRB4D3 complex together with a longer 80 bp dsRNA corroborates with the above observations (Fig S11A-S11C).

Based on titration of proteins with 40 bp RNA, we find that the apparent K_d_ of the interaction of DRB7.2, DRB4, and DRB7.2:DRB4 complex with 40 bp RNA is ∼ 2.2 μM, 3.6 μM, and 1.0 μM, respectively (Table 4). The higher apparent affinity of DRB7.2:DRB4 with pre-siRNA is probably due to the presence of multiple dsRBDs in the complex. Interestingly, 40 bp RNA interacts with both DRB7.2 and DRB4, and their complex with the Hill coefficient is more than 2 in all cases (Fig. 6E and Table 4). The Hill coefficient larger than 1 is often attributed to the intrinsic cooperativity of binding interaction; however, in our case, it implies that DRB4, DRB7.2, and their complex are probably interacting with the multiple consecutive minor-major-minor groove structure present on a 40 bp RNA.

**Table 4:** Apparent binding affinities (K_d_) and parameters derived from gel shift assays.

| SI No. | RNA | Protein | $K_d$ ($\mu\text{M}$)<sup>a</sup> | $n_H$ | $R^2$ |
| --- | --- | --- | --- | --- | --- |
| 1. | 40 bp RNA | DRB7.2 | $2.28 \pm 0.27$ | $2.68 \pm 0.77$ | 0.9839 |
| 2. | 40 bp RNA | DRB4 | $3.60 \pm 0.31$ | $2.44 \pm 0.45$ | 0.9915 |
| 3. | 40 bp RNA | DRB7.2:DRB4 | $1.02 \pm 0.17$ | $2.05 \pm 0.65$ | 0.9782 |

<sup>a</sup>Values derived from the average of three experiments.

Our data demonstrate that DRB7.2 dsRBD is capable of interacting with dsRNA of a varied length as well as with DRB4D3, through a distinct interaction interface. To validate if DRB4D3 and dsRNA occupy separate binding sites on DRB7.2 and do not interfere with individual recognition, we titrated the pre-formed complex of DRB7.2M:40 bp RNA with the increasing concentration of DRB4D3 (Fig. 6F). We observe that the preformed complex of DRB7.2M:40 bp RNA is not disrupted by DRB4D3 (Fig. 6F, similarly for 80 bp dsRNA (Fig. S11D)).

It is noteworthy that the oligomeric nature of DRB7.2M induces the formation of a mixture of the high order oligomeric complex of DRB7.2M eeven inn thee presence of dsRNA, which lags its migration (Fig. 6F). Whereas, at the equimolar and above equimolar concentrations of the DRB7.2M and DRB4D3, the monomeric nature of the heterodimer bound to the 40 bp RNA is compact in size, allowing it to migrate relatively faster (Fig. 6F).

The ITC titration of DRB4D3 with the preformed complex of DRB7.2M:80 bp RNA yielded a K_d_ ∼ 9.5 nM (Fig.S11E), comparable to the affinity measured for DRB7.2M:DRB4D3 in the absence of dsRNA. Overall, these results suggest that DRB7.2 can independently assemble heteromeric complex with DRB7.2M:DRB4D3 and DRB7.2M:dsRNA, and acts as a molecular interface for the formation of the ternary complex of dsRNA:DRB7.2M:DRB4D3 by sandwiching between dsRNA and DRB4.

Structural and biochemical studies presented reveal that DRB7.2 harbors two independent surfaces for simultaneous association with DRB4 and dsRNA. Besides, the ability of DRB7.2 to recognize dsRNA and that DRB4 dsRBDs bind to dsRNA ultimately leads to sequestering pre-siRNA by DRB7.2:DRB4 to stall the subsequent endonuclease-driven cleavage mediated by DCL3.

## DISCUSSION

The mechanism responsible for the competition to capture endoIR-precursor dsRNA by DCL3 versus DRB7.2:DRB4 complex remains elusive. As known from previous studies, many dsRBPs assist the corresponding Dicers to initiate and execute the RNAi pathway (Jouravleva *et al* 2022; Liu *et al* 2018; Wang *et al* 2021; Wilson *et al* 2015; Yamaguchi *et al* 2022), and therefore, the formation of the DRB7.2:DRB4 assembly to stall Dicer entry on the precursor dsRNA is quite contrasting.

Another important observation during the study was the conformational heterogeneity due to loosely coupled oligomeric states of the interaction domains of DRB7.2 and DRB4 in their solo form. The remarkable improvement in the NMR spectrum of the complex and the very high affinity of association affirm that the interaction domains attain a favorable heterodimeric state as a complex. Moreover, the evolution of DRB7.2 specific β2-β3 loop and the longer β3 remains a crucial step in the formation of the high affinity DRB7.2M:DRB4D3 complex as the key interactions for the complex formation are mediated by residues belonging to the extended β3 strands in DRB7.2M and the β1 strand of DRB4D3. Undoubtedly, it marks the first step in sequestering the endo-IR precursor RNA, as proposed in the model (Fig 7).

**Figure 7:**
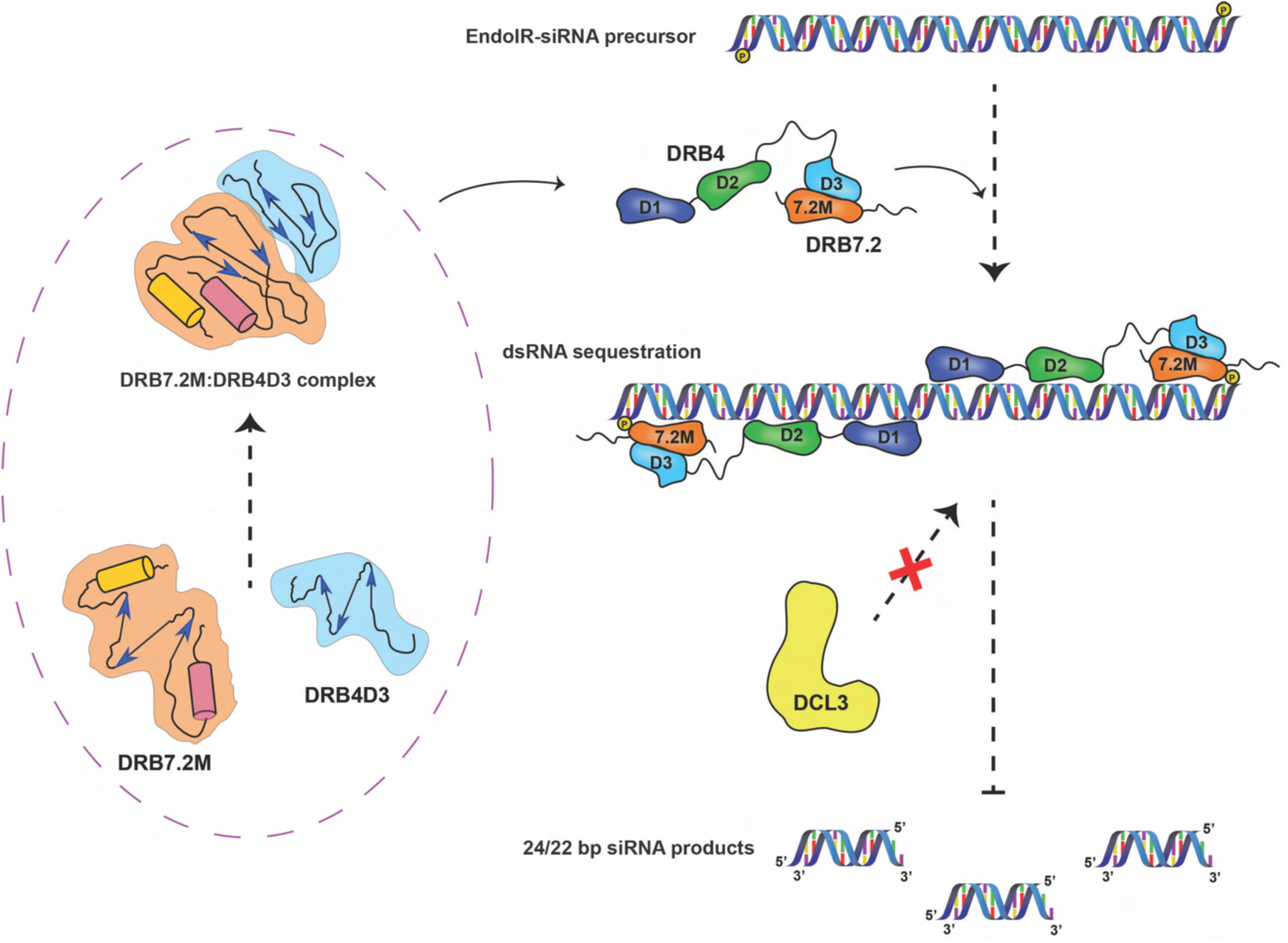
Proposed model for the dsRNA sequestration mechanism by DRB7.2 and DRB4 complex. The functional domains, DRB7.2M and DRB4D3 of DRB7.2 and DRB4, respectively, assemble a stable and very high-affinity heterodimeric complex such that DRB7.2M is accessible to the substrate RNA without steric hindrance. The ternary complex composed of DRB7.2, DRB4, and endo-IR precursor RNA is formed by the interaction of N-terminal dsRBDs of DRB4 and the sole dsRBD of DRB7.2 that can recognize the dsRNA fold. The formation of the DRB7.2:DRB4 complex increases the availability of the dsRBD pool that is necessary to prevent the access of DCL3 to the endo-IR precursor RNA, which leads to stalling of the production of 24/22 bp siRNA duplexes. The formation of the DRB4:DRB7.2 complex is a critical step in regulating the endo-IR precursor RNA processing. The structural divergence in DRB4D3 from its homologs in humans and flies (TRBPD3/R2D2D3) is a key event that allows plants to tailor the components of the RNAi machinery for diverse gene regulatory pathways.

The ability of DRB7.2 to simultaneously interact with substrate dsRNA and DRB4 using two independent binding sites is unprecedented. Unlike ribosomes, Dicers, and Argonautes, which typically recognize RNA and protein substrates through multiple domains, DRB7.2 achieves this dual recognition using two opposing surfaces within its dsRBD. This distinctive capability of the DRB7.2 dsRBD to concurrently recognize dsRNA and the DRB4D3 domain represents a significant and novel discovery highlighted in our study.

Earlier we have shown that the N-terminal DRB4 dsRBD1 is tailored to recognize the dsRNA and DRB4 dsRBD2 has ephemeral dsRNA binding activity in the presence of DRB4 dsRBD1 (Chiliveri *et a*l 2017). The binding study with 40 bp pre-siRNA presented here suggests that DRB4 dsRBDs and DRB7.2 dsRBD remain successful in populating multiple dsRBDs to sequester the pre-siRNA, thus potentially restricting access of pre-siRNA to DCL3 (Fig 7).

As evident from the transcript levels annotated at the bioanalytical resource for the plant biology (Toufighi K *et al*, 2005), DRB4 transcripts are twice more abundant than DRB7.2, which ensures that abundant DRB4 is ubiquitously available in the cytoplasm for binding with a diverse set of partners such as DCL4 and DRB7.2. As two independent processes involving DRB4, viz., the processing of the *trans*-acting siRNA by DCL4:DRB4 and sequestering endo-IR precursors by DRB7.2:DRB4 occur in the nucleus, plants have to devise an impeccable mechanism for DRB4 to choose between these two partners for efficient utility. We hypothesize that DRB4 interacts with either DRB7.2 or DCL4 post-translationally and the NLS on DRB7.2 and on nuclear localizing isoforms of DCL4 ensures DRB4’s journey to the nucleus. Therefore, by virtue of strong affinity, DRB4 gets tugged in the nucleus with its respective partners, DRB7.2 and DCL4. While the interaction interface of DCL4:DRB4 is not yet understood, the uniquely structured DRB4D3 primarily drives DRB7.2:DRB4 complex formation.

Furthermore, our results deliberate on the formation of DCL4:DRB4 mediated RNAi initiation complex formation in plants. In humans and flies, the RNAi initiation complex formation requires interaction of the C-terminal dsRBD or unstructured C-terminal region of dsRBP with the inter-helicase domain of corresponding Dicers. However, the DRB4D3 fold, despite being located at the C-terminus of DRB4, exhibits a very different structure from the canonical C-terminal α-β-β-β-α dsRBD of TRBP (*H. sapiens*) and R2D2 (*D. melanogaster*). Remarkably, DRB4D3 adopts a newly identified structural fold composed of three anti-parallel β strands arranged as β1-β3-β2. Despite extensive sequence and structural motif searches, no domain with such an abutted β strand arrangement has been identified during the course of this study. Moreover, following the formation of the DRB7.2M:DRB4D3 complex, DRB4D3 is left with only a little free surface for any additional interaction owing to its relatively small size compared to TRBPD3/R2D2D3. Therefore, DRB4 probably utilizes the remaining unstructured PxxP motif-rich C-terminus or the N-terminal dsRBDs to mediate the DCL4:DRB4 association. Alternatively, the fold attained by DRB4D3 upon binding with DRB7.2M may be an evolved feature enabling interactions with multiple partners, potentially adopting different conformations depending on the binding context.

Hence, our work implies that the organization of the plant RNAi initiation complex is distinct from its non-plant higher eukaryotes, as DRB4D3 is most likely impaired in directly associating with DCL4. Finally, we propose that the Dicer:dsRBP organization for dsRNA processing has evolved differently between plants and animals. Consequently, our results provide a foundation for future studies to unravel the precise organization of Dicer:dsRBP complexes across organisms possessing RNAi pathways and open up newer avenues for understanding the evolutionary diversification of RNA silencing mechanisms.

## METHODS

### Protein expression and purification

DRB4 (1-355) (Uniprot id: Q8H1D4), DRB4D1D2 (1-153), DRB4C (154-355), and DRB4D3 (294-355) constructs were cloned, expressed and purified as described (Chiliveri *et al*, 2017).

The genes encoding DRB7.2 (M1-V190) (Uniprot id: F4JHB3) and DRB7.2M (T71-G162) were PCR amplified from pSTM3-DRB7.2 plasmid and cloned into pETtrx-1b vector using gene-specific primers. Next, DRB7.2 and DRB7.2M were expressed in *E. coli* BL21 (RIPL) by inducing (OD_600_ ∼ 0.6) with 0.5 mM Isopropyl β-D-1-thiogalactopyranoside (IPTG) for 18 h at 18 °C. DRB7.2 and DRB7.2M were initially purified by affinity chromatography (His_6_ tag), followed by TEV protease cleavage and QFF anion-exchange chromatography. Subsequently, size exclusion chromatography was done using G75 Superdex preparative column. DRB7.2N (M1-Q81) was cloned in a pET28a vector and induced in *E. coli* BL21 (RIPL) with 0.1 mM IPTG and 3% ethanol at OD_600_ ∼ 0.6 at 15 °C for 24 h. DRB7.2N construct was purified by affinity chromatography (His_6_ tag), followed by size exclusion chromatography using G75 Superdex preparative column. ^15^N labeled samples were prepared by growing bacterial cells in an M9 minimal medium containing D_2_O and ^15^N NH_4_Cl as the sole nitrogen source. Additionally, ^13^C D-glucose was added to the above medium for preparing U-[^13^C, ^15^N], perdeuterated samples. Additional details of cloning, expression, and purifications are provided elsewhere (Paturi & Deshmukh, 2023).

### NMR spectroscopy and solution structure of DRB7.2M and DRB4D3

All NMR spectra were recorded on Bruker Avance Neo 600 MHz spectrometer, equipped with a triple-resonance cryogenically cooled TCI probe at 25 °C. TOPSPIN 4.0 (Bruker Biospin) and CARA-v.1.8.4 (Keller, 2004) were used for data processing and analysis, respectively.

An equimolar complex formed by U-[^13^C, ^15^N], perdeuterated DRB7.2M, and perdeuterated DRB4D3 was used to obtain the backbone chemical shift assignment of DRB7.2M. Similarly, perdeuterated, U-[^13^C, ^15^N] DRB4D3 and perdeuterated DRB7.2M were complexed at the equimolar ratio to obtain backbone chemical shift assignments of DRB4D3. For sidechain chemical shift assignment, the NMR data were obtained on a sample containing equimolar complex formed by partially ^2^H (∼35-40%), U-[^13^C, ^15^N] DRB7.2M, and perdeuterated DRB4D3 as well as partially ^2^H (∼35-40%), U-[^13^C, ^15^N] DRB4D3 and perdeuterated DRB7.2M. Backbone and sidechain chemical shifts of DRB7.2M and DRB4D3 were adapted from BMRB accession no. 51790 and 51791, respectively (Paturi & Deshmukh, 2023).

^1^H-^1^H distance restraints were obtained using 3D NOESY-^15^N HSQC (*τ*_m_ = 80 ms and 150 ms) and 3D NOESY-^13^C HSQC (*τ*_m_ = 80 ms) on the complex formed by ^2^H (∼50%), U-[^13^C, ^15^N] DRB7.2M:perdeuterated DRB4D3 and ^2^H (∼35-40%), U-[^13^C, ^15^N] DRB4D3:perdeuterated DRB7.2M. NOE cross peak intensities were used to calculate distances and were grouped into the following categories: strong (1.8-3.0 Å), medium (1.8-4.0 Å), weak (1.8-5.0 Å), and very weak (1.8-6.0 Å). Dihedral angle values were derived from the backbone chemical shift values (H^N^, N, C’, C_α_, C_β_) using TALOS-N (Shen *et al*, 2013).

The initial structures of DRB7.2M were calculated using an extended polypeptide chain as a template and the dihedral and distance restraints NOEs under Cartesian dynamics simulated annealing protocol in Xplor-NIH (Schwieters *et al*, 2003). A square-well penalty function is used for all restraints with the following force constants: 50 kcal mol^-1^ Å^-2^ and 200 kcal mol^-1^ rad^-2^ for NOEs and torsional angles, respectively. Furthermore, the ten lowest energy structures were refined using ROSETTA fixed backbone refinement protocol using experimental restraints (Ramelot *et al*, 2009). The final ensemble of the ten lowest energy structures is used for further analysis and has been deposited in the PDB with code 8Z49.

We used ab-initio relax protocol in Rosetta data (Raman *et al*, 2010) to calculate the solution structure of DRB4D3 using the chemical shift and sparse amide-amide NOE data. In a nutshell, the PSI-BLAST program (Altschul *et al*, 1997) is executed for five iterations to generate a position specific scoring matrix (PSSM). Furthermore, TALOS-N (Shen *et al*, 2013) prediction was used for the generation of the chemical shift table to predict structural factors, such as backbone torsion angles and secondary structures. Later, the chemical shifts, TALOS-N prediction, and PSSM were used as inputs for the ROSETTA fragment-picking procedure to generate two sets of *de novo* protein fragments, i.e., 3-mer and 9-mer. ^1^H-^1^H NOEs sampled for both proteins were further used as distance restraints. The Rosetta prediction calculations were run for 5000 structures after eliminating the N-terminus unstructured region encompassing E294-S316 to avoid erroneous prediction of structural elements in the disordered region. The acceptance of Rosetta-generated models is assessed by plotting the convergence of Rosetta total energy against Cα RMSD (Fig. S4A). The ensemble of the ten lowest energy structures of DRB4D3 is available in the PDB databank with code 8Z4A.

All the RMSD were calculated using the RMSD trajectory tool built in the VMD software (Humphrey *et al*, 1996) where all the secondary structure residues were fitted to an average structure.

### Crystallization and structure determination of DRB7.2M:DRB4D3 complex

The equimolar ratio of DRB7.2M and DRB4D3 was mixed, and the complex was purified by size-exclusion chromatography using a Superdex 75 preparative column in potassium phosphate buffer (50 mM potassium phosphate buffer (pH 7.0), 50 mM NaCl, 50 mM Na_2_SO_4_, 2 mM DTT).

Peak fractions from the SEC, which correspond to DRB7.2M:DRB4D3 in concentrations ranging from 2 mg/mL to 8 mg/mL and a drop ratio of 1:1 (protein:well solution), were used to set up sitting-drop vapor diffusion experiments. Sparse matrix screens such as Index HT screen and Crystal Screen HT in 96-well crystallization plates were used to identify crystallization conditions for the DRB7.2M:DRB4D3 complex. The crystallization plates were incubated at 293 K. Diamond-shaped tetragonal crystals were visible after 3 days in crystallization conditions containing 0.15 M sodium citrate tribasic dihydrate (pH 5.6), 20% v/v 2-propanol, and 20% w/v polyethylene glycol 4,000. Further, the expansion screen was set up with the purified complex concentrated at 6 mg/ml and subjected to crystallization at 293 K with the hanging-drop vapor diffusion technique. The crystallization drops contained 2 μL protein solution mixed with 2 μL of reservoir solution. The DRB7.2M: DRB4D3 complex crystals were directly vitrified in the liquid nitrogen stream without any cryoprotectant. Diffraction data were collected at 100 K on an in-house X-ray diffractometer (Rigaku Micromax 007 HF with a Mar345-dtb image plate detector) using Cu Kα (*λ* = 1.54179 Å). The X-ray diffraction images were indexed and integrated using the processing routine available in the CCP4 software suite (Project, 1994).

The crystals belong to the space group *I* 4, with unit-cell parameters a = 93.94, b = 93.94, and c = 102.81 Å. The crystal structure of DRB7.2M:DRB4D3 was solved by molecular replacement method using PHENIX (Adams *et al*, 2010) with the homology models of DRB7.2M and DRB4D3 obtained from AlphaFold colab (Jumper *et al*, 2021). Iterative model building and refinement were performed using COOT (Emsley & Cowtan, 2004) and the PHENIX (Adams *et al*, 2010), which resulted in a final R_cryst_ and R_free_ of 20.20% and 25.34%, respectively. Residues T71-R82 of DRB7.2M and E294-K305 of DRB4D3 could not be modeled owing to poor electron density in these regions. We performed the buried surface area calculations of DRB7.2M:DRB4D3 complex using PISA (Krissinel & Henrick, 2007). The surface plot of DRB7.2M:DRB4D3 implies that it assumes an axially symmetrical ellipsoid shape with an overall molecular dimension of 36.6 x 39.7 x 52.8 Å. The data collection and refinement statistics are summarized in Table 3, and the PDB ID obtained is 8IGD for the DRB7.2M:DRB4D3 complex.

### Isothermal titration calorimetry (ITC)

ITC experiments were performed on an automated MicroCal PEAK-ITC or MicroCal ITC-200 at 25 °C with stirring (750 rpm). All samples were degassed at room temperature before use. Protein and RNA samples were prepared in potassium phosphate buffer (50 mM potassium phosphate (pH 7.0), 50 mM NaCl, 50 mM Na_2_SO_4_, 2 mM DTT). Titrants (200 μM DRB4 and 200 μM DRB4D3) (Syringe volume: 40 μL) were titrated with the first injection of 0.4 μL followed by 18 injections of 2 μL (150 s intervals) into titrates (20 μM DRB7.2 and 20 μM DRB7.2M) (Cell volume: 280 μL). MicroCal Origin 7.0 was used to analyze the data and create the ITC plots. After baseline integration, the data were zeroed based on the assumption that the final injections represent the heat of dilution.

### Electrophoretic mobility gel shift assay

In the present work, we used electrophoretic mobility gel shift assays (EMSA) with 40 bp pre-siRNA and 80 bp RNA with 1 nt 3’ overhang. As DRB7.2 is known to preferentially bind with longer dsRNA with a higher affinity (Montavon *et al*, 2017), we performed EMSA of various constructs used in this study with 40 bp RNA and 80 bp RNA. Additionally, we estimated the K_d_ of DRB7.2, DRB4, and DRB7.2:DRB4 complex with commercially synthesized 40 bp RNA, i.e., pre-siRNA with 1 nt 3’ overhang, which is a known DCL3 substrate (Wang *et al*, 2021), (guide: 5’ P-AACAAGCGAAUGAGUCAUUCAUCCUAAGUCUGCAUAAAGU 3’, complementary: 5’ ACUUUAUGCAGACUUAGGAUGAAUGACUCAUUCGCUUGUUC 3’), using gel mobility shift assays.

For each assay, 20 μL reaction containing 750 nM dsRNA and varying protein concentrations were incubated for 30 min at room temperature in buffer containing 50 mM potassium phosphate (pH 7.0), 50 mM NaCl, 50 mM Na_2_SO_4_, and 2 mM DTT.

The protein:RNA complex was separated on a 7% native PAGE, and the electrophoresis was performed at a constant 80 V for 2 h, RT. The gels were visualized using SYBR Gold Nucleic Acid Gel Stain, the bound and unbound RNA intensities were calculated from the gels using ImageJ software, and the triplicate data (fraction bound vs protein concentration) was plotted to calculate the apparent K_d_ and Hill coefficient using non-linear regression curve fit analysis (Growth/Sigmoidal using Hill equation) using OriginPro 8.0.

## ACCESSION NUMBER

The structures determined are deposited in the PDB data bank under accession code 8Z49 (Xplor-NIH with Rosetta refined NMR structure of DRB7.2M), 8Z4A (Rosetta derived NMR structure of DRB4D3), and 8IGD (crystal structure of the DRB7.2M:DRB4D3 complex).

## Extended Data

Extended data containing NMR relaxation studies, a table, and eleven figures are available.

## ACKNOWLEDGEMENTS

We acknowledge Prof. Patrice Dunoyer (Institut de Biologie Moléculaire des Plantes du CNRS, Strasbourg) for the generous gift of recombinant DRB7.2 plasmid. We acknowledge Sayali Khisty and Dr. Sai Chaitanya Chiliveri for their general assistance during various stages of the project. Staff at the X-ray facility and Dr. R. Sankaranarayanan are acknowledged for their assistance related to the diffraction data collection and general discussion during the crystal structure determination. The central instrumentation facility of CCMB is acknowledged for maintaining facilities and equipment. For generous access to NMRbox, we acknowledge the National Center for Biomolecular NMR Data Processing and Analysis, a Biomedical Technology Research Resource (BTRR).

## AUTHOR CONTRIBUTIONS

MVD planned the study and designed experiments. SP, DP, PB, RA, NW, and MVD performed experiments and interpreted the results. PB, NW, and SP performed the gel-shift assays. SP, DP, RA and NW assisted in ITC and NMR titration data collection. SP and DP performed the solution structure calculation of DRB7.2M and DRB4D3. SP performed the crystallization and structure refinement where KP extended assistance in the final stages of the refinement. MVD, DP, and SP wrote the manuscript.

## Extended data

The extended data contains NMR-driven ^15^N backbone relaxation studies, one table describing dynamics parameters derived from relaxation studies, and eleven supporting figures to the main text.

### 15N NMR backbone relaxation and model-free analysis

R_1_, R_2,_ and ^15^N-{^1^H} heteronuclear NOE experiments were performed on equimolar complexes of [U-^15^N] DRB7.2M:perdeuterated DRB4D3 and [U-^15^N] DRB4D3:perdeuterated DRB7.2M, where the concentration of each construct was 200 μM. Relaxation rates were estimated by fitting the peak intensities as a function of relaxation delay to a single exponential function and errors in the relaxation rates were calculated by running 500 Monte Carlo simulations using Relax (Bieri et al, 2011). The ratio of peak intensities with and without proton pre-saturation was used to estimate steady-state NOE, and the standard deviation in the NOE values was calculated from the RMS value of the background noise. R2R1_TM, Quadric diffusion (Lee et al., 1997), and the heterodimeric complex crystal structure were used to make an initial guess of the molecular rotational diffusion tensor from the R_2_/R_1_ ratios derived from the individual residues. Model-free analysis was performed with Model-free 4.1 (Mandel et al., 1995) interfaced with FAST-Modelfree (Cole et al., 2003). In addition, we have extensively used tools hosted by NMRbox for the relaxation data analysis (Maciejewski et al., 2017).

The ^15^N backbone relaxation rates (R_1_ and R_2_), the R_2_/R_1_ ratio, and the ^15^N-{^1^H}-heteronuclear NOE (hetNOE) as a function of residues is represented in Fig. S6 for DRB7.2M:DRB4D3 complex. We have excluded residues T71, T72, E73, D127, S128, K144, and A146 of DRB7.2M and residues E294-C298, K306, K307, and Q321 of DRB4D3 from the analysis due to exchange broadening as manifested by the poor exponential decay fits during data analysis. Residues M124 and K143 of DRB7.2M were excluded due to a lack of backbone chemical shift assignments. The average R_1_, R_2_, and hetNOE values obtained for the structured region of DRB7.2M are 0.8 s^-1^, 20 s^-1,^ and 0.8, respectively, whereas, the DRB4D3 correspondingly yielded R_1_, R_2_, and hetNOE values of 0.8 s^-1^, 22 s^-1^, and 0.7, respectively. While the residues T75-S84, C111, H112, K125, and T131 in DRB7.2M display flexibility as they assume a hetNOE value of 0.3 or lower, the β2-β3 region appears to be rigid. Similarly, in DRB4D3, the N-terminal region comprising V299-I309 is also flexible, as seen from the average hetNOE values of 0.3 or lower.

We obtained an initial estimation of the molecular rotational diffusion tensor using the R_2_/R_1_ ratio for residues that are not affected by fast ps-ns timescale internal dynamics or by slow μs-ms timescale dynamics. Using this analysis, we find that the average *τ*_c_ of the complex is 15.27±0.65 ns, considering only the mean R_2_ and R_1_ (Table S1). Additionally, using the quadric diffusion (Lee et al., 1997), we have estimated the initial values of the principal components, the parallel (D_ǁ_ = D_ZZ_) and perpendicular (D_⊥_ = 0.5 (D_XX_+D_YY_)), of the diffusion tensor. The F-test value analysis suggested that the rotational diffusion of the complex is best described by an axially symmetric diffusion tensor over a fully anisotropic tensor. However, the estimated rotational correlation time and the diffusion anisotropy (D_ratio_ = D_ǁ_/D_⊥_) are highly identical for both models. Considering an axially symmetric tensor, we obtained an average *τ*_c_ = 14.43±0.34 ns with D_ratio_ = 1.25±0.23 for the complex (Table S1). Simultaneously, we obtained a *τ*_c_ value of 14.09 ns using the Hydrodynamic radius approach embedded in the program HYDRONMR (Torre et al., 2000). Furthermore, we have optimized the diffusion tensor using the ^15^N backbone relaxation data through extended model-free formalism (using the crystal structure of the complex as the input structure) to derive residue-specific amplitude and fast timescale motions (ps-ns). Iterative model-free fitting was set up with the estimates derived from the above approaches, where the diffusion tensor along with the parameters for internal motions S^2^ and *τ*_e_ is optimized to fit the R_1_, R_2_, and NOE data according to the model selection procedure described by Mandel et al. (Mandel et al., 1995). After convergence, we obtained optimized values for *τ*_c_ = 15.79±0.05 ns and diffusion anisotropy as D_ratio_ = 1.13±0.05. NMR relaxation analysis resulted in the rotational correlation time (*τ*_c_) of 15.8 ns, which corresponds to a molecular weight of ∼ 25 kDa of 1 unit of DRB7.2M:DRB4D3 complex.

The other features derived from the model-free analysis, such as the square of the order parameter (S^2^), model selection, fast-timescale internal motion (*τ*_e_), and the conformational exchange rate (R_ex_) as well as the atomic displacement parameter (B-factor) derived from the crystal structure are as shown in Fig. S7. During the final selection of the dynamical model for the individual residues, we observed that the majority of the DRB7.2M:DRB4D3 complex residues assume an average value of 0.85 for the S^2^ indicating that the majority of the complex behaves as a rigid entity with no significant dynamics. Nonetheless, the residues C111, H112, and T131 of DRB7.2M and residues L308, I309, and W328 of DRB4D3 show a drop in S^2^ below a value of 0.5 suggesting that these regions might possess local flexibility. Unsurprisingly, the decrease in the S^2^ values corroborates with an increase in the crystallographic B-factor, as shown in Fig. S7.

As expected, the majority of the residues within the secondary structural elements of the complex assume Model1, where motions on a slow time scale are negligible, and the generalized order parameter is represented by the fast timescale motions only. In DRB7.2M, residues selected for Model2 are Y90, L92, H97, E108, G109, M113, G118, V132, L133, N138, and G162 with *τ*_e_ ranging between 40-140 ps, whereas S84, which is sandwiched between α0-α1, assumes a higher *τ*_e_ of 567 ps. Similarly, in DRB4D3, Model2 is chosen for G311, H314, L315, S316, T319, and G320 with relatively faster liberational motion represented by *τ*_e_ ranging from 750 ps to 1200 ps, while residues R326, E331, and D337 show *τ*_e_ < 500 ps. The selection of just a single residue in Model3 (D344 in DRB4D3, with R_ex_ = 7.126 s^-1^) and Model4 (K125 in DRB7.2M with *τ*_e_ = 71.2 ps and R_ex_ = 8.75 s^-1^) supports the fact that DRB7.2M and DRB4D3 attain a stable high-affinity complex devoid of slow exchange dynamics. Model5 is selected for C111, H112, K114, R129, I130, T131, K142, and K159 in DRB7.2M (*τ*_e_ ∼ 780-1200 ps) and L308, I309, M310, T312, I317, W328, T333, L334, V353, and K354 in DRB4D3 (*τ*_e_ ∼ 900-1850 ps), where 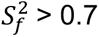 except T131 and W328.

**Table S1:** Diffusion tensor analysis for DRB7.2M:DRB4D3 complex based on the ^15^N backbone relaxation data.

| | Tensor | $\tau_c(\text{ns})^a$ | $D_{\text{ratio}}^b$ | $\chi^2$ | F |
| --- | --- | --- | --- | --- | --- |
| Estimated by R2R1_TM<sup>c</sup> | - | 15.27±0.65 | - | - | - |
| Estimated Diffusion tensor from R<sub>2</sub>/R<sub>1</sub> using quadric diffusion<sup>c</sup> | Isotropic | 14.36±0.04 | - | 6049 | - |
|  | Axial symmetric | 14.43±0.34 | 1.25±0.23 | 5875 | 1.04 |
|  | Fully anisotropic | 14.36±0.04 | 1.37±0.02 | 5913 | 0.11 |
| HYDRONMR<sup>d</sup> | - | 14.09 | 1.30 | - | - |
| Optimized<sup>e</sup> |  | 15.79±0.05 | 1.13±0.05 | - | - |

<sup>a</sup>Rotational correlation time obtained from the relation $\tau_c = (6D_{\text{iso}})^{-1}$ .
<sup>b</sup>Ratio of the elements of the diffusion tensor: for axially symmetric case: $D_{\text{ratio}} = D_{\parallel}/D_{\perp}$ and for the fully anisotropic case and hydrodynamic calculations $D_{\text{ratio}} = 2D_{\text{ZZ}}/(D_{\text{XX}} + D_{\text{YY}})$ .
<sup>c</sup>R2R1\_TM and quadric diffusion were used to evaluate residue-specific dynamic properties (Lee et al., 1997).
<sup>d</sup>Results from hydrodynamic calculations by the program HydroNMR (Ortega et al., 2005; Torre et al., 2000).
<sup>e</sup>Optimized results using model-free results analysis.

**Figure S1:**
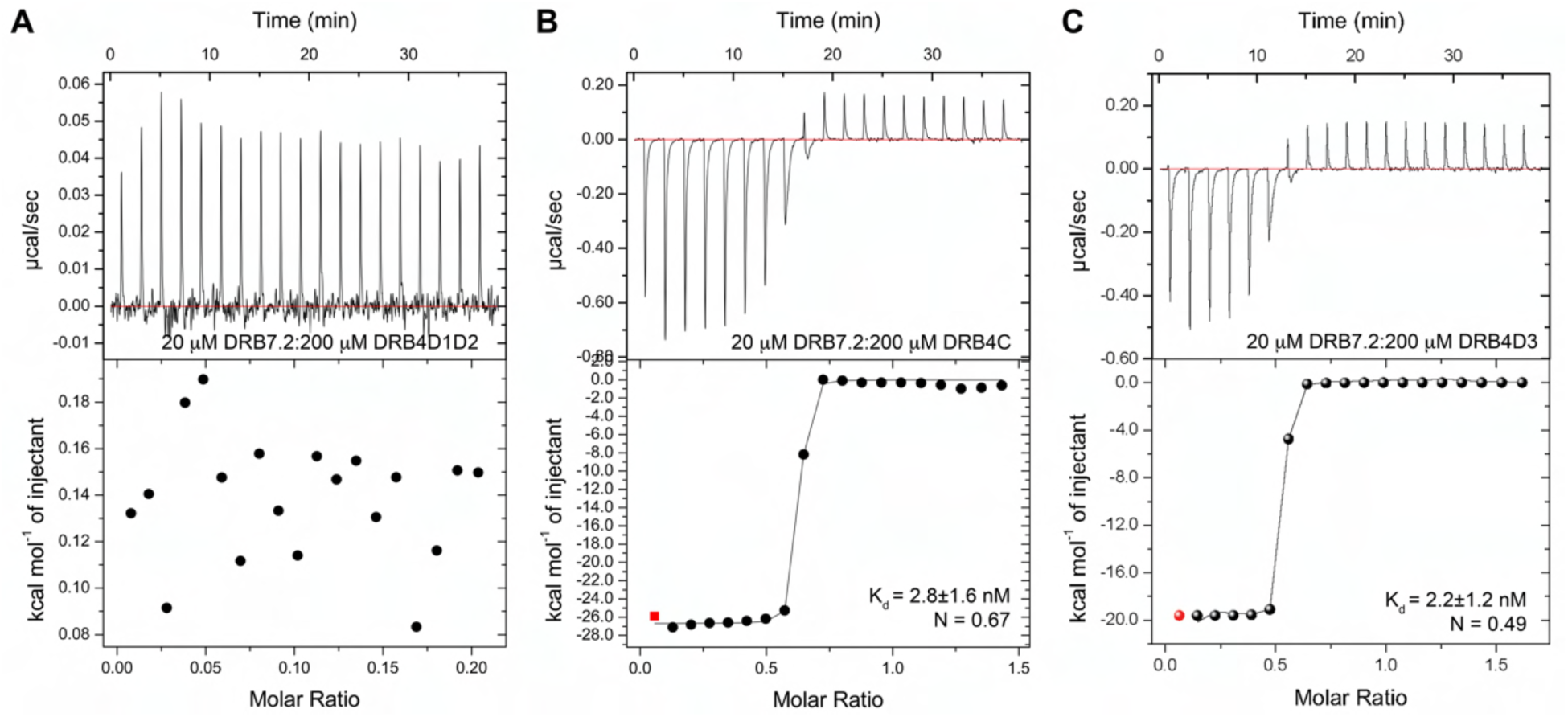
Interaction of DRB7.2 with DRB4D1D2, DRB4C, and DRB4D3. (A) ITC plot of DRB7.2 titrated with DRB4D1D2 shows no interaction, dictating the role of two N-terminal dsRBDs of DRB4 in dsRNA-specific interaction. (B) The binding isotherm of DRB7.2 with DRB4C shows an affinity of 2.8±1.6 nM, suggesting that both N-terminal DRB4 dsRBDs do not take part in the binding with DRB7.2. (C) The ITC plot of DRB7.2 with DRB4D3 showing an affinity of 2.2±1.2 nM suggests that DRB4D3 is necessary and sufficient to interact with DRB7.2.

**Fig S2.**
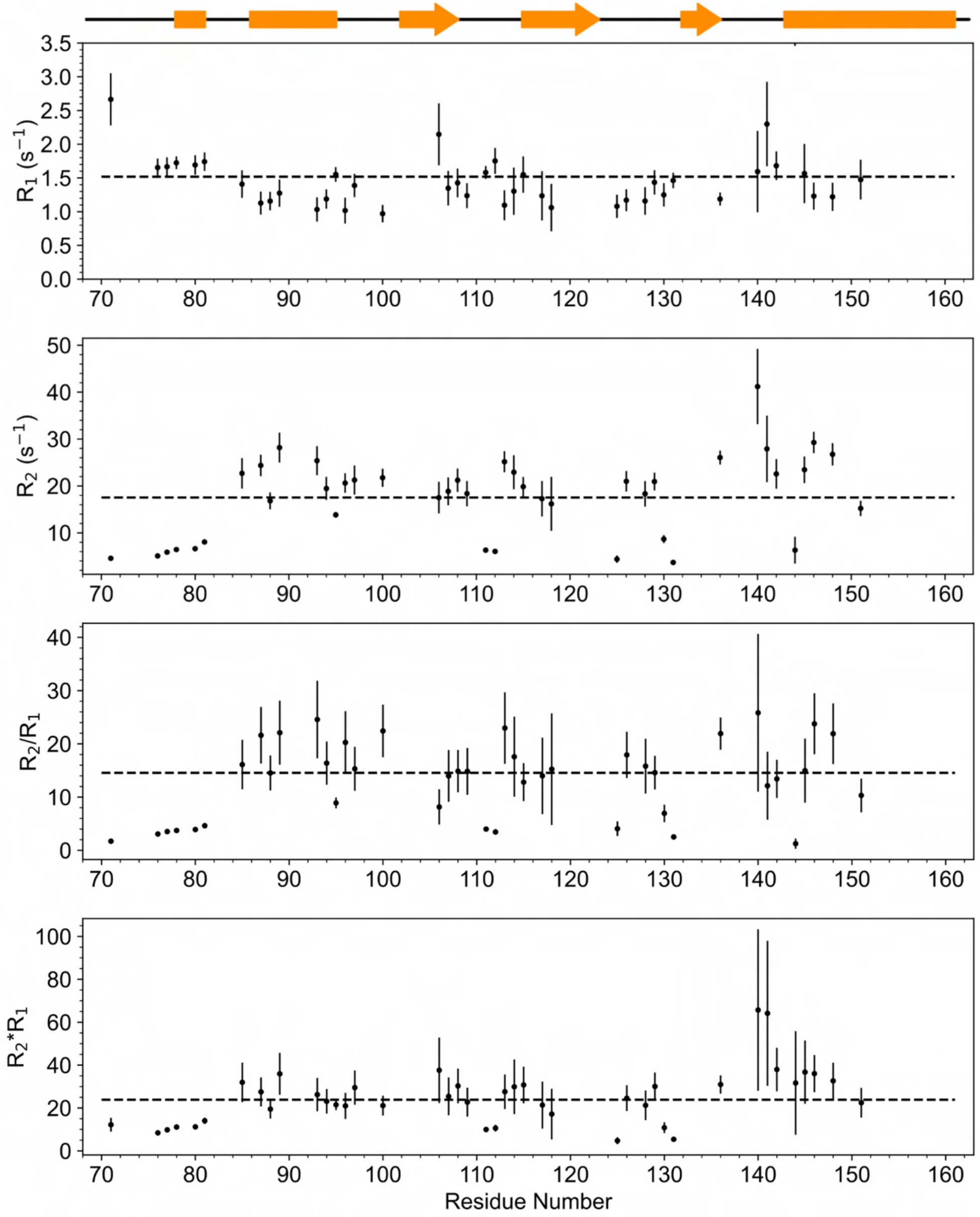
^15^N backbone relaxation studies and analysis on DRB7.2M. ^15^N R_1_, R_2_, R_2_/R_1,_ and R_2_*R_1_ were estimated unambiguously after transferring backbone chemical shift of 42 residues. The average R_2_/R_1_ corresponds to 14.56, whereas R_2_*R_1_ appears to be 23.84. The estimated *τ*_c_ from the data corresponds to 11.89 ns, whereas the theoretical *τ*_c_ for a monomeric 10 kDa globular protein should be ∼ 5-6 ns. The enhanced *τ*_c_ in the case of the DRB7.2M implies that DRB7.2M is probably experiencing slow to intermediate exchange between monomer:dimer and higher-order oligomeric states.

**Figure S3:**
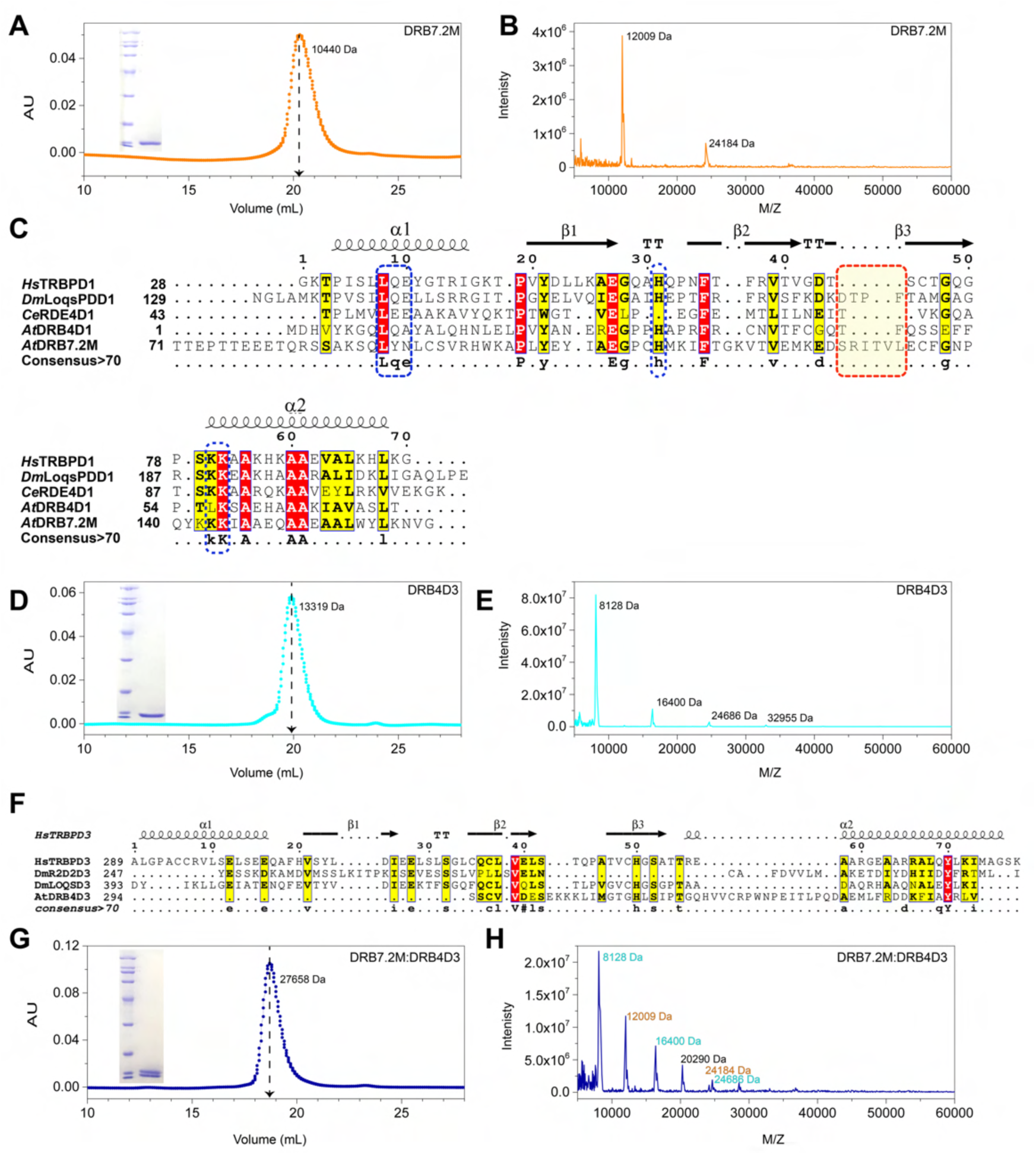
Characterization and primary sequence analysis of DRB7.2M and DRB4D3. (A) Size exclusion chromatography (SEC) profile of DRB7.2M corresponding to the molecular weight of 10440 Da, (B) MALDI-TOF MS of DRB7.2M resulted in 12009 Da as observed MW. A small peak appearing at 24184 Da is most likely a dimeric fraction, implying DRB7.2M may remain in dynamic equilibrium of monomer:dimer in the free state. (C) Sequence alignment of DRB7.2M with homologous dsRNA binding domains. The dsRNA binding residues involved in making a tripartite contact with dsRNA are highlighted with blue dashed boxes. The extended β2-β3 loop specific to DRB7.2M is marked with red dashed boxes. (D) Size exclusion chromatography profile yields 13319 Da as MW of DRB4D3, whereas (E) corresponds to the MALDI-TOF MS profile of DRB4D3 in the free state. Multiple fragments at higher order oligomeric state in DRB4D3 suggest its dynamic equilibrium of monomer to higher order oligomers (up to tetramer or so) in the free form. (F) The sequence comparison of DRB4D3 with *Hs*TRBPD3, *Dm*R2D2D3, and *Dm*LoqsPBD3 implies that the C-terminal region of DRB4 is divergently evolved from its non-plant higher eukaryotes homologs. (G) Size exclusion chromatography profile of DRB7.2M:DRB4D3 complex gives an MW of 27658 Da. (H) The corresponding MALDI-TOF MS shows the major monomeric peaks corresponding to the complex but also small populations of oligomeric states of individual proteins. The analytical SEC data was collected on 1 mg/mL protein concentration on a G75 Superdex analytical column, and the molecular weights were calculated using standards (Cytiva). MALDI-TOF analysis was performed with 0.5 mg/mL, 2 mg/mL, and 5 mg/mL concentrations. SDS PAGE for samples post-SEC is given as an inset in the corresponding panels. Secondary structural elements, α-helices and β-strands derived from the PDB structures of *Hs*TRBPD1 (PDB ID: 5N8M) and *Hs*TRBPD3 (PDB ID: 4WYQ) in panels C and F, respectively, are represented at the top of the alignment. Highlighted in yellow are the invariant residues, and in red are conserved by >70%.

**Figure S4:**
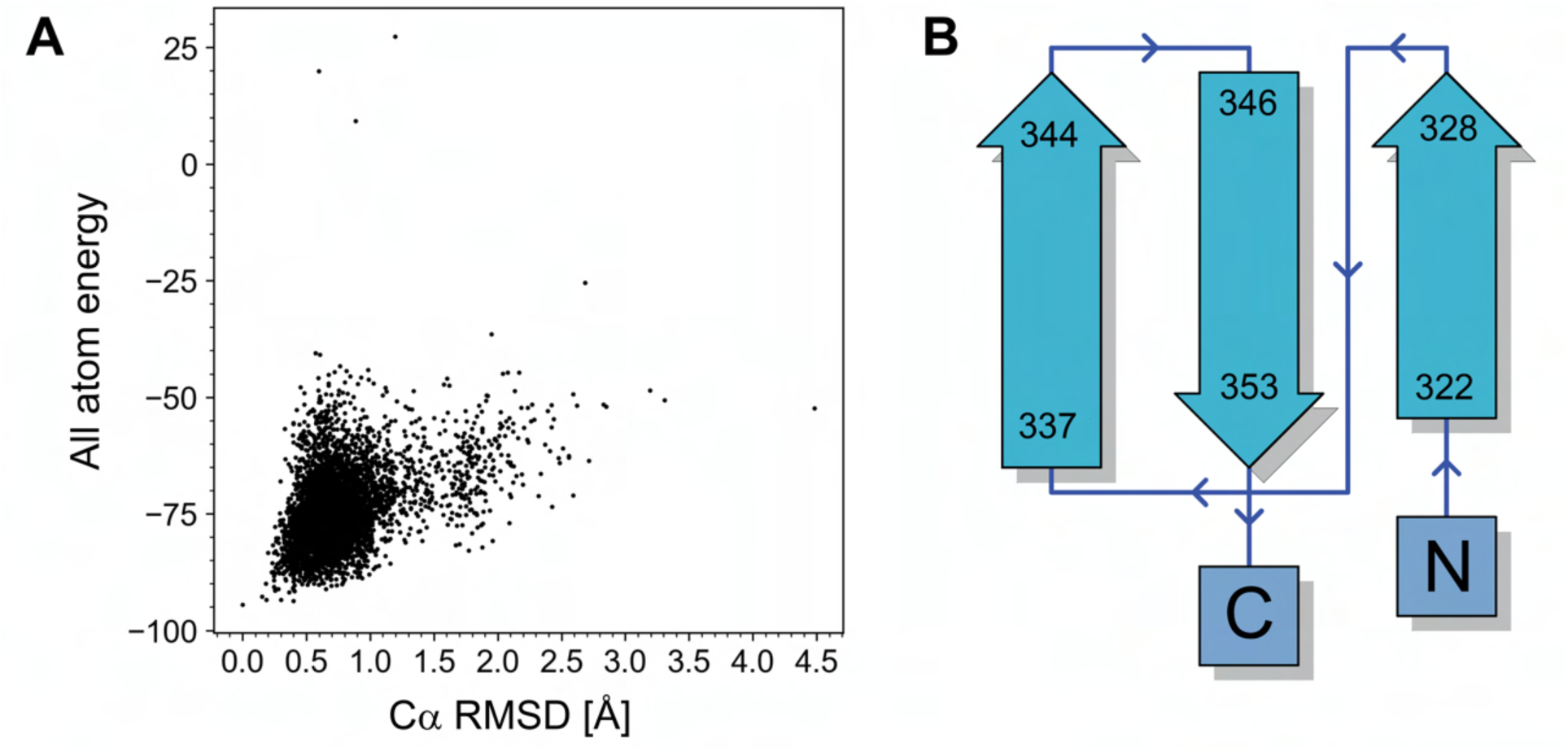
Rosetta convergence plot and topology map for DRB4D3. (A) Cα RMSD vs. all-atom energy in DRB4D3 (convergence at 5000 structures). The RMSD was calculated against the lowest energy structure. (B) The topology of the novel fold adopted by DRB4D3 upon binding with DRB7.2M is derived from the PDBsum1 server (Laskowski et al. 2022).

**Figure S5:**
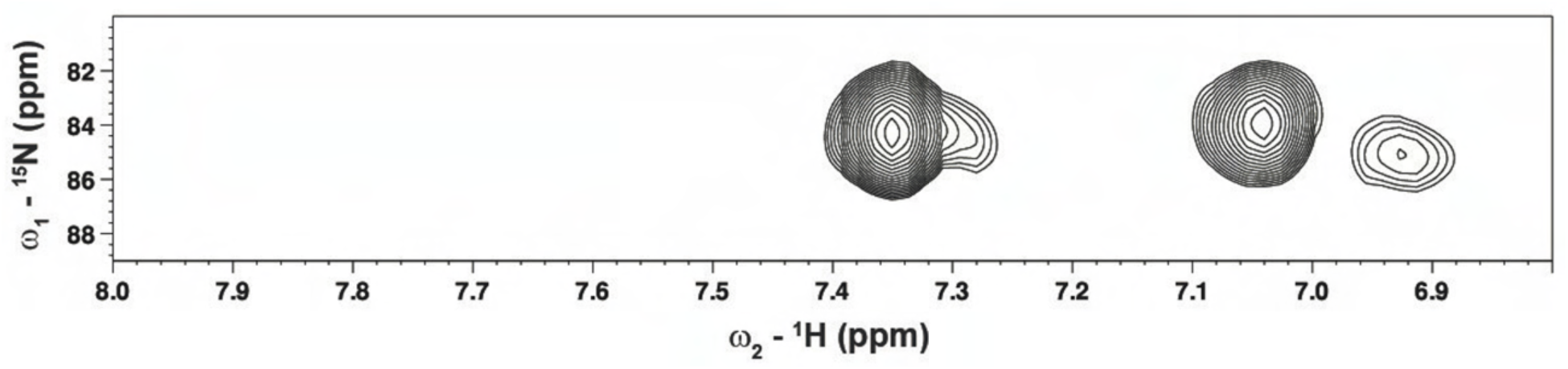
Salt bridge interaction in DRB7.2M:DRB4D3 complex. ^1^H-^15^N HSQC-TROSY spectrum of DRB4D3 showing the region of Arginine guanidino group (Nε-Hε) amide protons of the sidechain involved in salt-bridge formation through either intra or intermolecular interaction. For DRB7.2M, the low ^15^N frequencies (82-86 ppm) peaks in the ^1^H-^15^N HSQC-TROSY were not observed, suggesting that the salt-bridge interactions involve the amide sidechains in DRB4D3.

**Figure S6:**
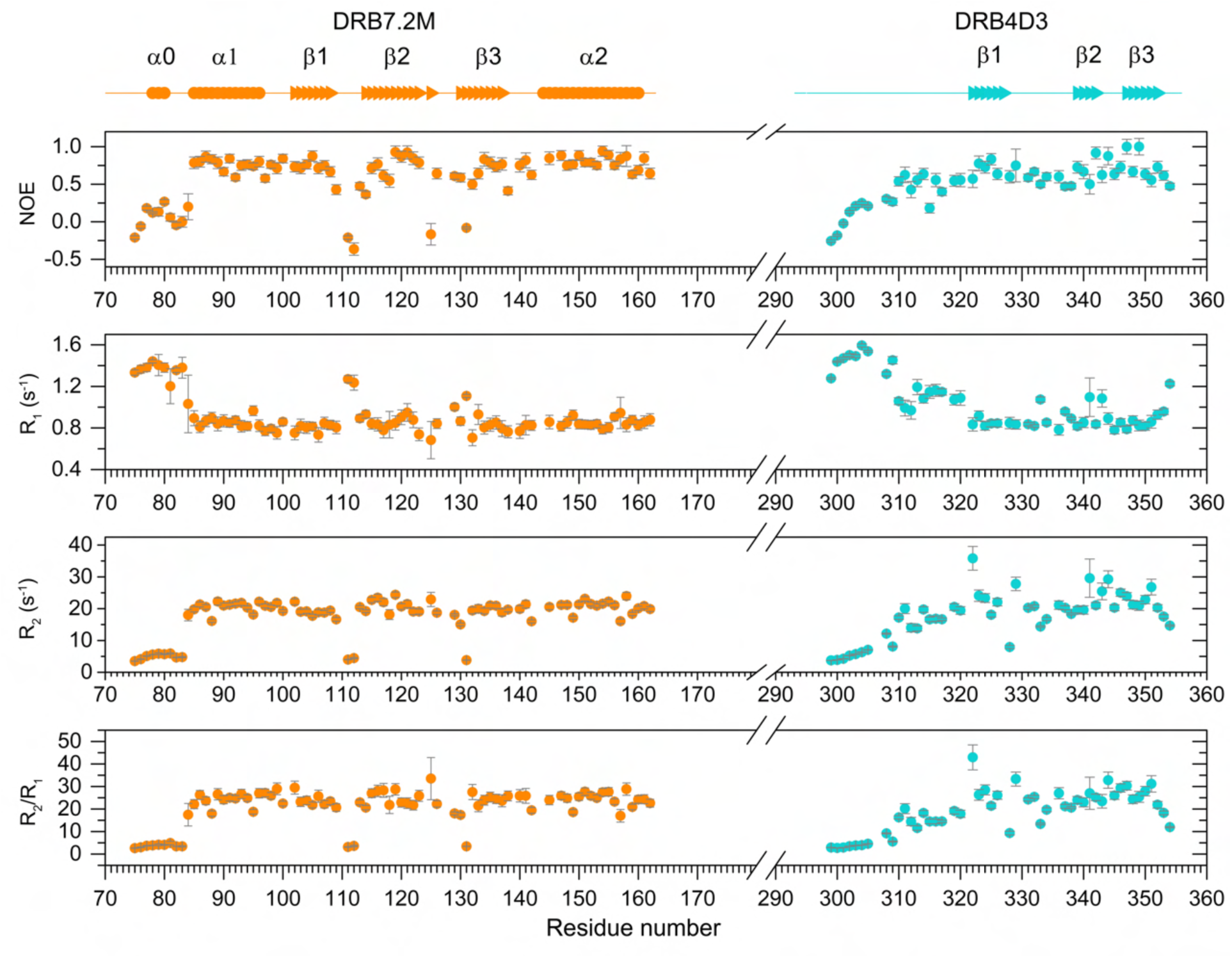
^15^N backbone relaxation studies of DRB7.2M and DRB4D3 complex. ^15^N-{^1^H} hetero-nuclear NOE, R_1_, R_2_, and R_2_/R_1_ obtained at 600 MHz are represented as a function of residue number. Secondary structural elements, α-helices (circles) and β-strands (arrows), are represented on the top. The data obtained for the N-terminal region of DRB7.2M and DRB4D3 was not used further for the model-free analysis as these regions were missing in the crystal structure.

**Figure S7:**
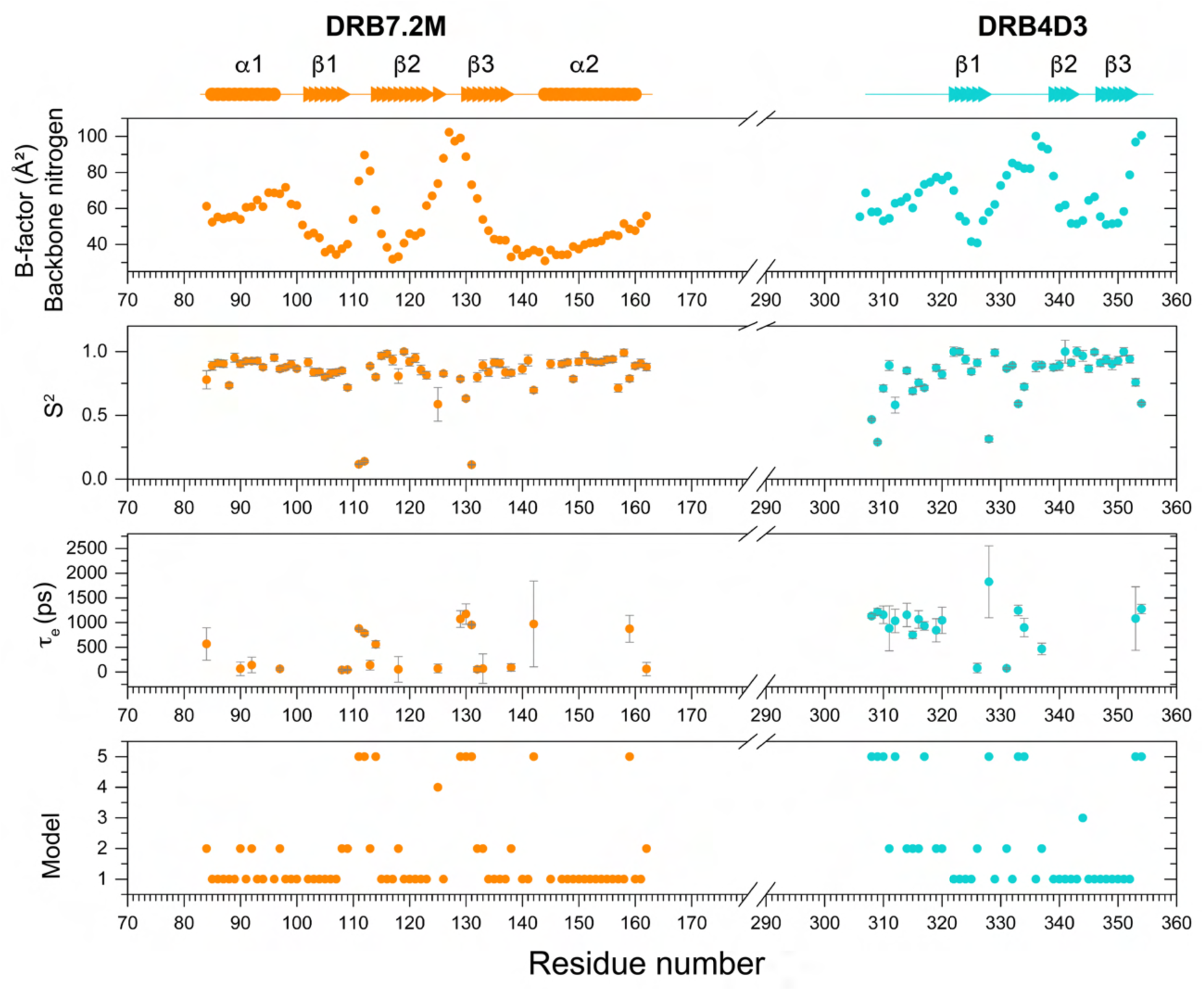
Backbone dynamics studies for the heterodimer complex formed by DRB7.2M:DRB4D3. The B-factors derived from the electron density map for backbone nitrogens obtained from the structure of the complex, the square of the order parameter (S^2^), the fast-timescale internal motion time (*τ*_e_), and the dynamic model selection using Model-free formalism as described by Mandel et al. (Mandel et al., 1995) as a function of residue number. Secondary structures are annotated as filled circles and filled arrows for α-helices and β-strands, respectively. The dynamic information derived from the NMR relaxation and x-ray electron density map suggests that both interacting domains are overall rigid in the complex and do not exhibit significant internal motions at the interaction interface. However, the selection of Model2/Model5 for E108, G109, M113, H112, and K142 coupled with the inherent resonance broadening in K143 and K144 imply that the dsRNA binding site experiences dynamics at a fast to intermediate timescale (ns-ms).

**Figure S8:**
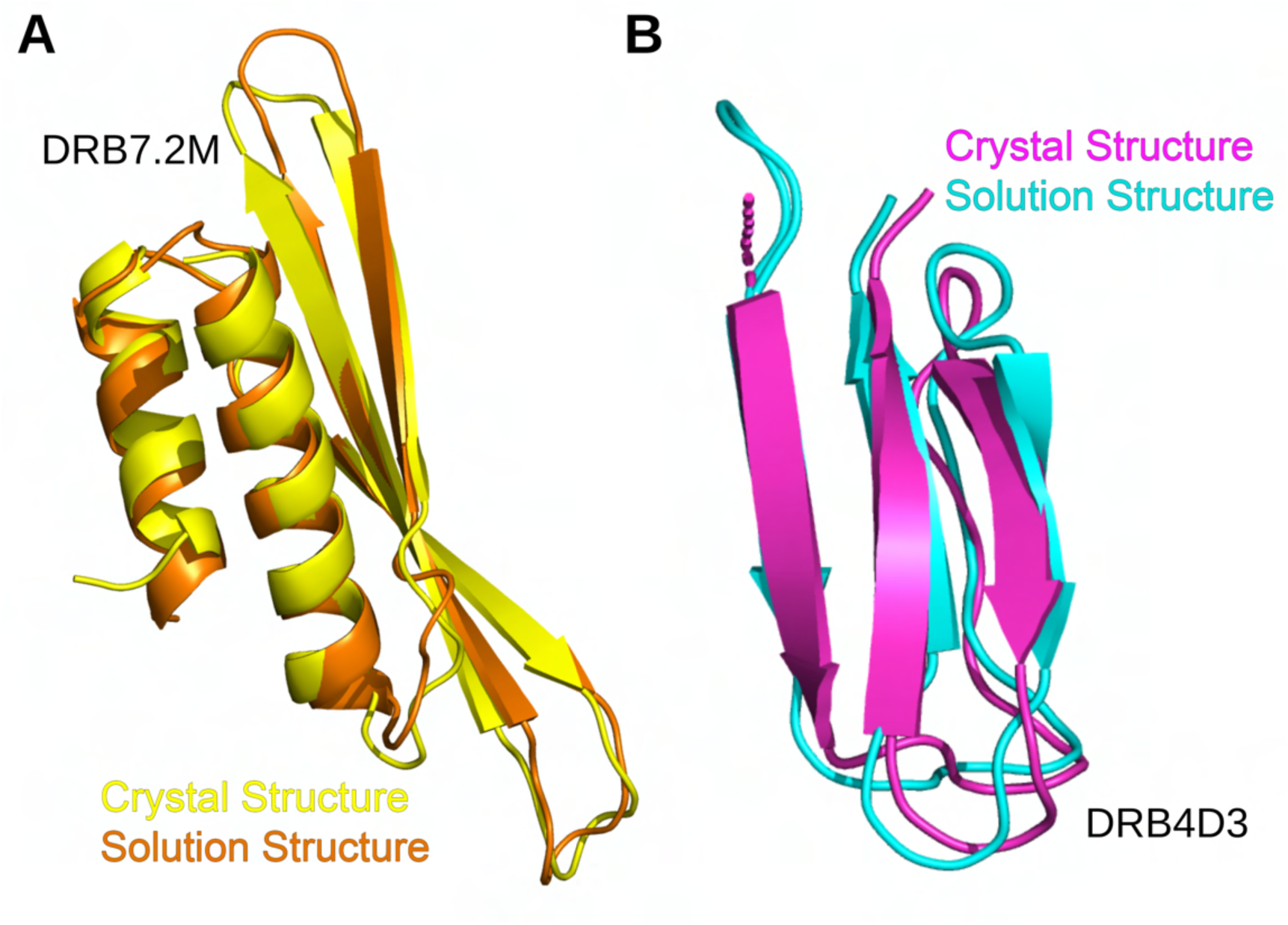
Comparison of the solution and crystal structures of DRB7.2M and DRB4D3. (A) Overlay of the lowest energy structure obtained using Xplor-NIH and crystal structure of DRB7.2M with backbone RMSD of 0.87 Å shows a similar arrangement of the dsRBD fold apart from a minute deviation in the length of β2 and β3 strands. The density for residues T71-R82 is absent in the crystal structure mainly due to the inherent flexibility of the region, as mentioned above. (B) Overlay of lowest energy structure obtained using Rosetta and X-ray structure of DRB4D3 with backbone RMSD of 0.59 Å with a similar arrangement.

**Figure S9.**
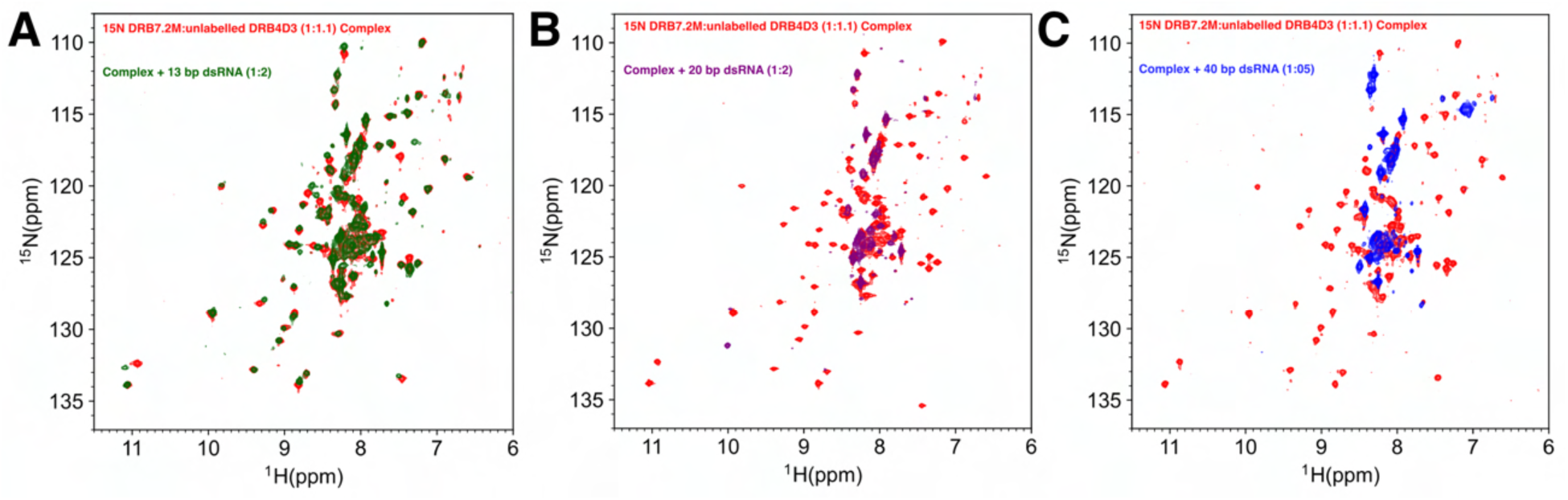
A full 2D ^15^N-^1^H HSQC of DRB7.2M:DRB4D3 complex with different dsRNA. (A) The overlay of the ^1^H-^15^N HSQC of 150 μM ^15^N DRB7.2M precomplexed with 165 μM unlabeled DRB4D3 (red) and upon titration with 300 μM 20 bp dsRNA (green). (B) The overlay of the ^1^H-^15^N HSQC of 150 μM ^15^N DRB7.2M precomplexed with 165 μM unlabeled DRB4D3 (red) with 300 μM 20bp siRNA duplex (magenta). (C) The overlay of the ^1^H-^15^N HSQC of 150 μM ^15^N DRB7.2M precomplexed with 165 μM unlabeled DRB4D3 (red) with 75 μM 40bp dsRNA (blue).

**Figure S10:**
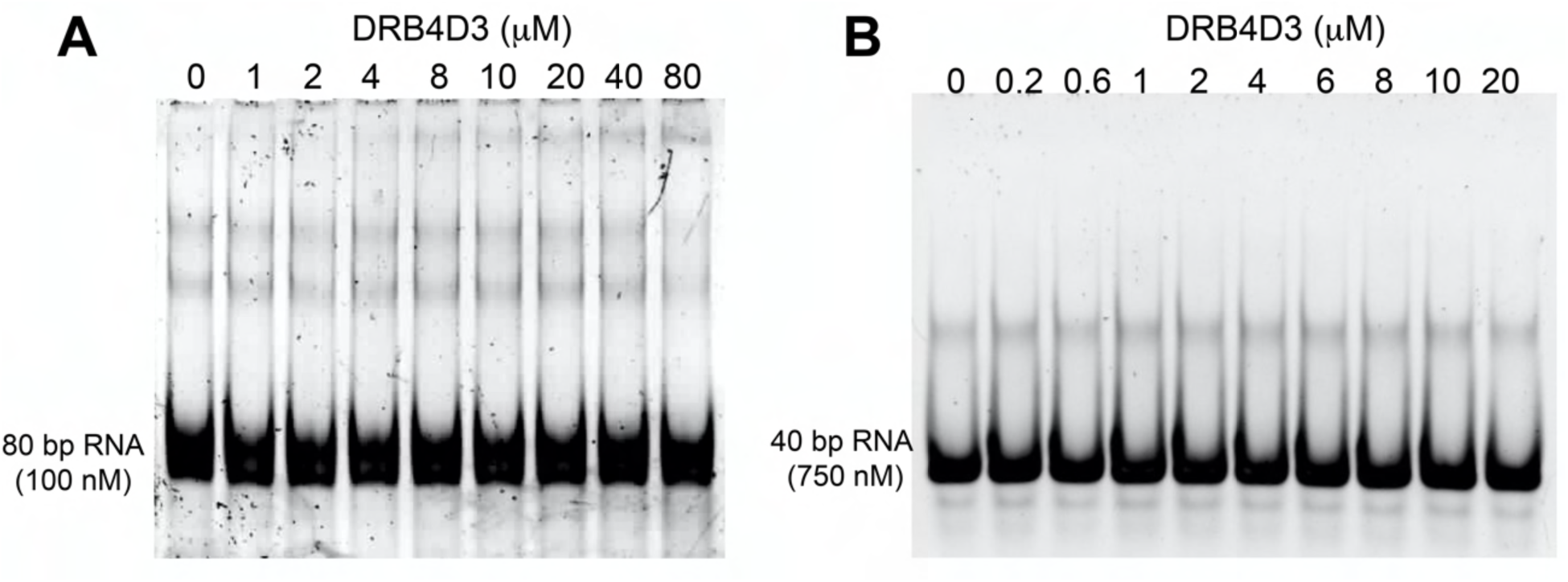
Gel shift assays for DRB4D3. EMSA of DRB4D3 with (A) 80 bp RNA and (B) 40 bp RNA, where samples were separated on a 7% native PAGE. Subsequently, electrophoresis was performed at 80 V for 3 h at 4 °C, and the gel was visualized with SYBR Gold staining. The data shows that DRB4D3 does not bind to dsRNA.

**Figure S11:**
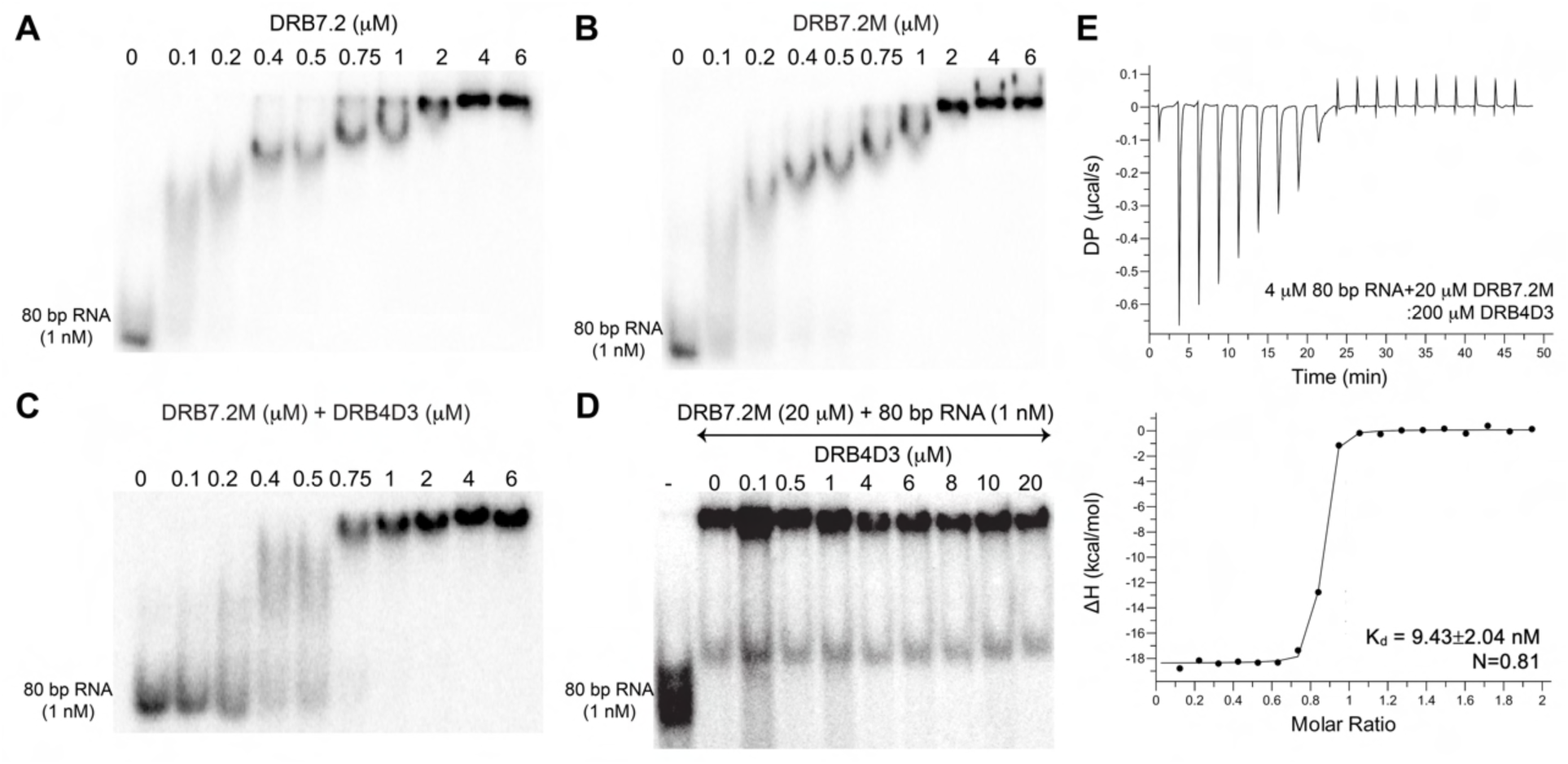
Functional characterization of DRB7.2 with 80 bp blunt end dsRNA. Gel shift assay of (A) DRB7.2, (B) DRB7.2M, and (C) DRB7.2M:DRB4D3 complex with internally labeled 80 bp dsRNA. Eighty nucleotide long sense and antisense RNA (sense: 5’ GGGUGCUGUUUCUCGUGUUCGUGUUCGUUUCUCUUCUCUUGUCCUUGUUCUGUUCUCCUUUGUUCGUUCCUGUUCCCCUU 3’, and antisense: 5’ AAGGGGAACAGGAAC GAACAAAGGAGAACAGAACAAGGACAAGAGAAGAGAAACGAACACGAACACGAGAAACAGCACCC 3’) were internally labeled with [α-^32^P] CTP using *in vitro* transcription and purified. The titration in panel A-C with increasing concentration of dsRNA shows saturation of protein at around ∼ 2 μM concentration. For the gel shift assays, recombinant proteins (∼0.1-10 μM) were incubated with [α-^32^P] labeled 1 nM 80 bp dsRNA for 1 h at 4 °C in buffer containing 50 mM potassium phosphate (pH 7.0), 50 mM NaCl, 50 mM Na_2_SO_4_, 2 mM DTT. Samples were separated on a 7% native PAGE. Subsequently, electrophoresis was performed at 80 V for 3 h at 4 °C, and the gel was analyzed using autoradiography. (D) Gel shift assay of a preformed complex of DRB7.2M:80 bp dsRNA with increasing concentrations of DRB4D3. (E) ITC-derived titration study of DRB7.2M:80 bp dsRNA complex with DRB4D3. The K_d_ corresponding to 9.43±2.04 nM with N=0.81 was obtained by fitting the isotherm to one site binding mode. The data implies that the interactions between DRB7.2M, dsRNA, and DRB4D3 occur independently, and the ternary complex formation by the three components transpires through exclusive binding events.

